# Quantifying spatiotemporal dynamics and noise in absolute microbiota abundances using replicate sampling

**DOI:** 10.1101/310649

**Authors:** Brian W. Ji, Ravi U. Sheth, Purushottam D. Dixit, Harris H. Wang, Dennis Vitkup

## Abstract

Metagenomic sequencing has enabled detailed investigation of diverse microbial communities, but understanding their spatiotemporal dynamics remains an important challenge. Here we present DIVERS, a widely applicable method based on replicate sampling and spike-in sequencing that quantifies the contributions of temporal dynamics, spatial sampling variability and technical noise to the variances and covariances of absolute bacterial abundances. Using high resolution time series profiling, we apply DIVERS to the human gut microbiome. Our method reveals complex spatiotemporal dynamics of individual gut bacteria and unmasks key features of their behavior hidden from previous analyses.

---

Metagenomic sequencing is widely used to explore patterns of bacterial abundances and the spectrum of functions carried out by diverse microbial communities^1–4^. However, as research efforts move beyond static descriptions of communities towards understanding their complex spatiotemporal dynamics^5–8^, a number of key challenges remain to be addressed. First, robust and quantitative frameworks are required to characterize both the temporal variability and spatial heterogeneity of bacterial abundances across environments^9,10^. Second, the effects of technical noise arising from sample preparation and sequencing must be quantified and accounted for^11,12^. Third, measurements of absolute bacterial abundances are necessary to correct for possible compositional artifacts associated with relative abundances^13,14^. Importantly, quantitative approaches for understanding both the spatiotemporal and noise profiles of microbial communities must be experimentally tractable, minimizing the number of sample preparations and costs associated with data collection, processing and sequencing.

To address these challenges, we have developed Decomposition of Variance Using Replicate Sampling (DIVERS), a broadly applicable method for metagenomic sequencing studies. DIVERS utilizes the laws of total variance and covariance to provide a principled mathematical approach for separating the contributions of time, spatial sampling location and technical noise to measured abundance variances for individual taxa and covariances for pairs of taxa:

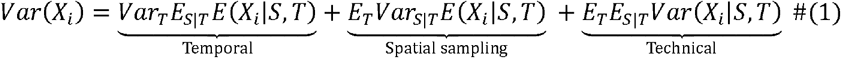

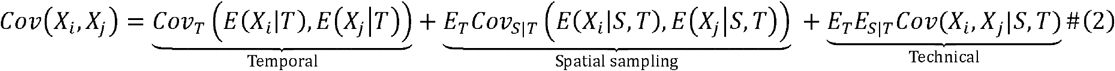

where *X*_i_ denotes the absolute abundance of an individual bacterial taxon *i*, *S* and *T* are space and time-associated random variables capturing the respective spatial and temporal processes affecting the abundance of taxon *i*, and *E*, *Var* and *Cov* denote the expectation, variance and covariance of random variables respectively.

We derived unbiased statistical estimators for each of the six terms in equations (1) and (2), and devised a workflow to enable their calculation directly using experimental measurements (**Supplementary Note**, Online Methods). Importantly, DIVERS requires only two samples obtained from randomly chosen spatial locations at each time point of a longitudinal microbiome study. One of these spatial replicates is then split in half to obtain two technical replicates, and absolute abundance measurements on the resulting three samples are performed using a spike-in procedure during sample processing^15,16^ (**Fig. 1a**, **Supplementary Note**, Online Methods). The key idea behind this approach is that bacterial taxa that exhibit genuine temporal fluctuations should also exhibit large abundance covariances between pairs of spatial replicates across time points. Spatial variability, quantified by differences in abundances between the two random locations, and technical noise result in decreased covariances. Interestingly, our sampling scheme and underlying mathematical model are conceptually similar to the dual reporter approach previously used to separate intrinsic and extrinsic sources of noise in gene expression profiles^17,18^ (**Supplementary Note**).

**Figure 1.**
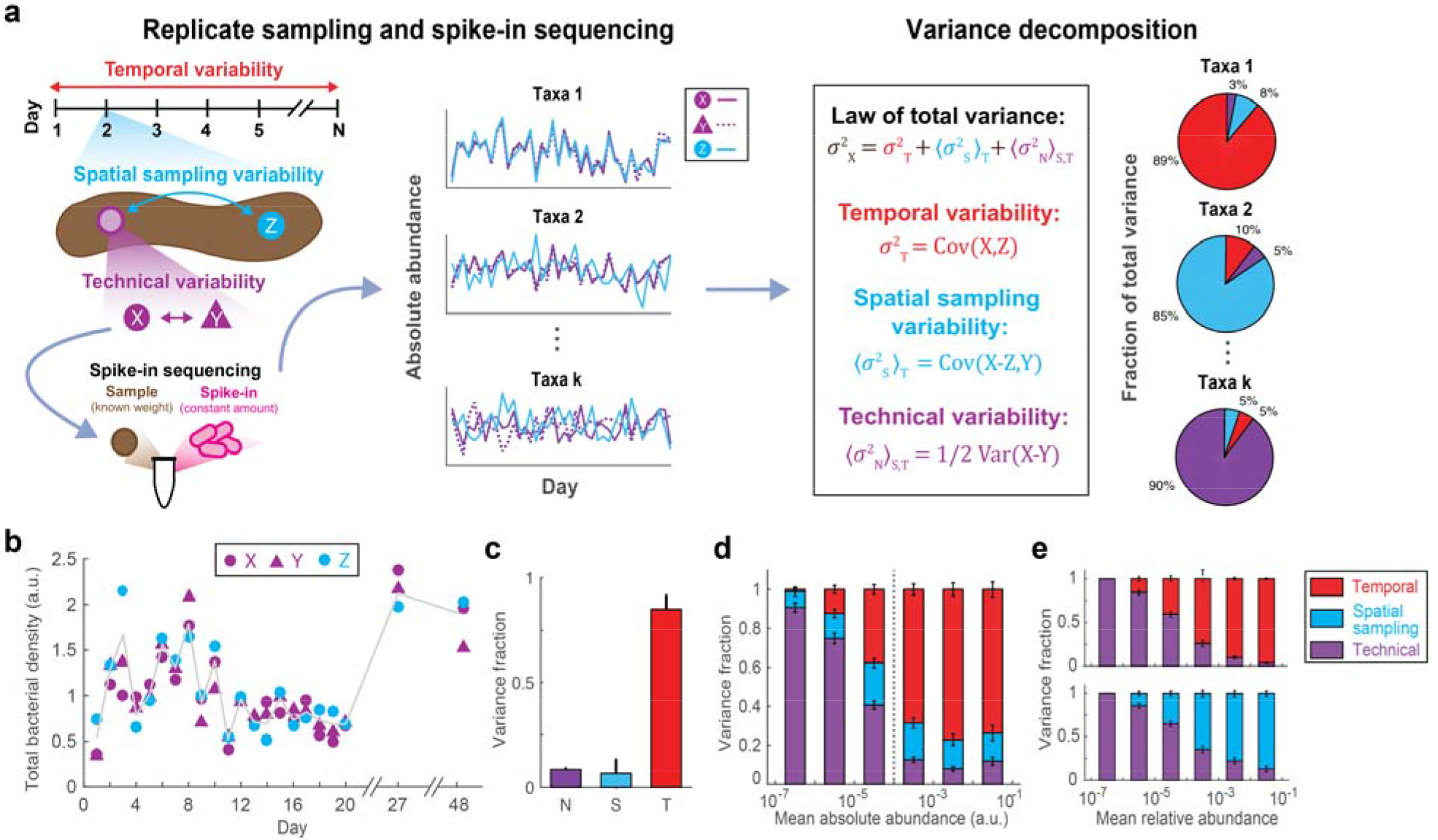
Variance decomposition of gut bacterial abundances using DIVERS. (**a**) Illustration of the DIVERS workflow applied to the fecal microbiome. Samples are collected from two random spatial locations (X/Y and Z, as shown on the left side of the figure) on each day of microbiome sampling, and two technical replicates (X and Y) are prepared from one these spatial locations. The resulting three samples from each day (X, Y and Z) are subjected to a custom spike-in procedure to estimate absolute bacterial abundances. The DIVERS variance decomposition model is then applied (right side of the figure) to abundance profiles of each taxa to quantify contributions of temporal variability, spatial sampling heterogeneity and technical noise to total abundance variability. (**b**) Temporal profiles of total bacterial densities in the human gut microbiome. X and Y correspond to technical replicate measurements of total bacterial density from a single spatial location, while Z corresponds to a second spatial replicate. Gray line shows the average of spatial replicates. Total bacterial densities are reported in arbitrary units and normalized to an average of one (Online Methods). (**c**) Variance fraction of total bacterial densities attributed to technical (N, purple), spatial sampling (S, blue) and temporal (T, red) factors as calculated by the variance decomposition model. (**d**) Variance decomposition of individual OTU abundances. Absolute OTU abundances were obtained by multiplying relative abundance profiles by the total bacterial density in each sample and are reported in arbitrary units (Online Methods). OTUs are binned by their mean abundance across all samples, and stacked bars show the average variance contribution of technical, spatial sampling and temporal sources to OTUs within each bin. Dashed vertical line corresponds to a mean absolute abundance of 10^−4^. Error bars represent the SEM. (**e**) Variance decomposition of microbiota abundances from control experiments. Top: DIVERS applied to stool samples without spatial variability; bottom: DIVERS applied to stool samples without temporal variability.

To demonstrate the ability of DIVERS to separate different sources of microbiota variability, we focused on the human gut microbiome, an ecosystem known to exhibit complex spatiotemporal dynamics^9,19^. We carried out 16S rRNA sequencing of fecal samples collected daily over the course of three weeks from a healthy male individual. (**Fig. 1a**, Online Methods). Using the obtained data, we first characterized total baseline bacterial abundance variation in the human gut microbiome. Consistent with previous results^13^, we found that total bacterial abundances fluctuated substantially across different samples (coefficient of variation = ~0.5) (**Fig. 1b**). Notably, the observed variability was dominated by daily temporal changes, with total bacterial loads remaining relatively constant across different spatial locations on each day (**Fig. 1c**).

Using measurements of total bacterial loads, we calculated the absolute abundances of all operational taxonomic units (OTUs) and then used DIVERS to decompose the abundance variance of individual OTUs (Online Methods and **Supplementary Note**). Interestingly, variance profiles exhibited two regimes when OTUs were grouped by average abundance, with a transition occurring at ~0.01% in relative abundances (**Fig. 1d** and **Supplementary Fig. 1**). Fluctuations of OTUs below this abundance cutoff could be primarily explained by technical sources of variability, generally consistent with Poissonian sampling noise (**Supplementary Fig. 2b** and **Supplementary Fig. 3**). In contrast, variability of OTUs above this cutoff largely reflected temporal changes (**Fig. 1d** and **Supplementary Fig. 2a**). Differences across spatial sampling locations also contributed a substantial fraction of total variability (on average ~20% across OTUs with an average absolute abundance > 10^−4^), demonstrating significant spatial heterogeneity of fecal samples.

**Figure 2.**
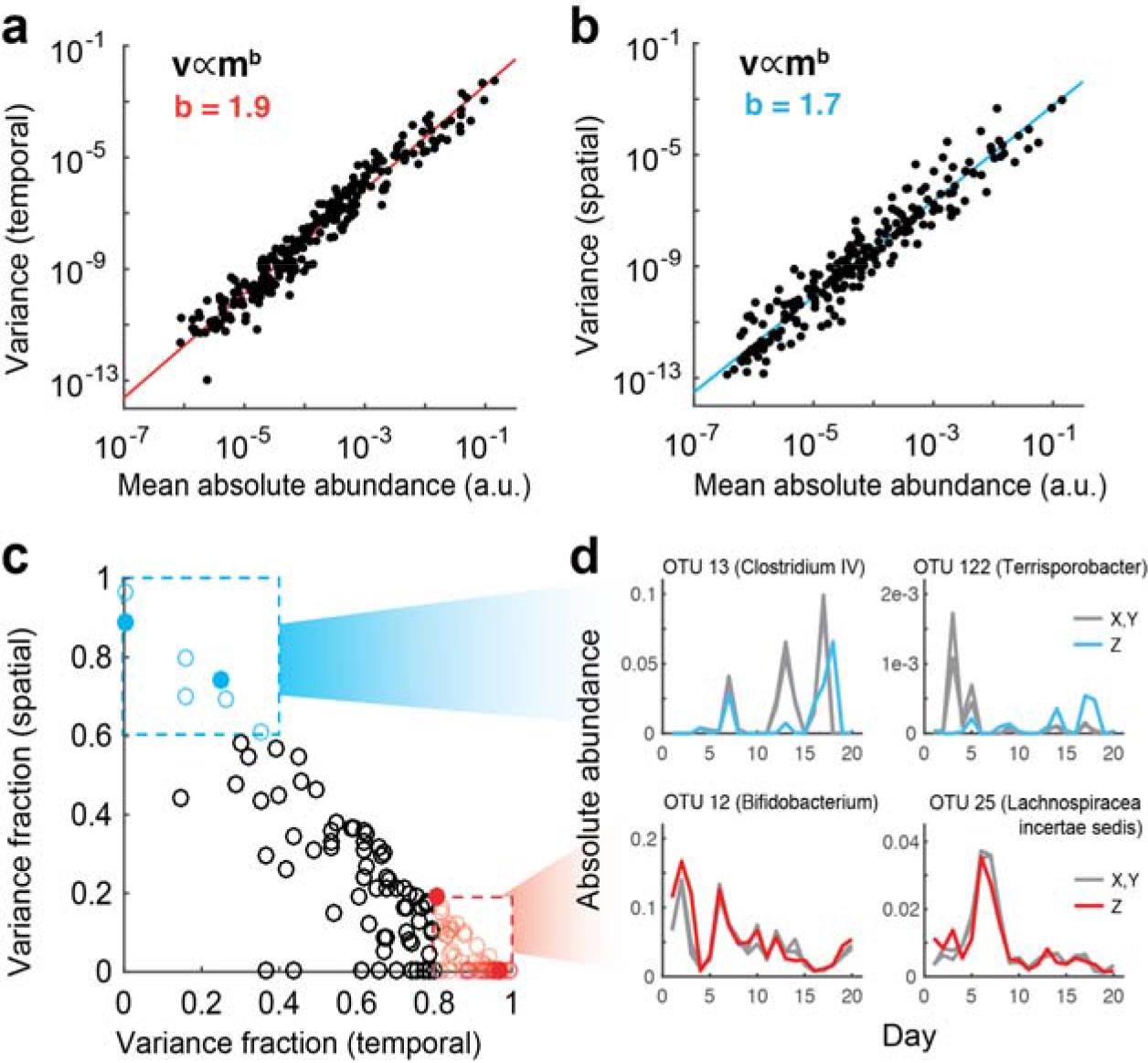
Temporal and spatial variation of human gut microbiota. (**a,b**) Temporal and spatial variances of human gut microbiota abundances calculated using the DIVERS variance decomposition model. Each data point corresponds to the average abundance (*m*) and the temporal (in **a**) and spatial (in **b**) abundance variance (*v*) of a particular OTU. Variances follow a power law of the form *v* ∝ *m*^*b*^ with exponents b=1.9 (temporal) and b=1.7 (spatial). (**c**) Identification of specific OTUs with either high temporal or high spatial variance contributions. Boxes indicate OTUs with a predominant contribution of temporal (variance fraction > 0.8, red) or spatial variances (variance fraction > 0.6, blue). Only abundant OTUs (mean absolute abundance > 10^−4^) are shown. (**d**) Time series of individual OTUs, corresponding to filled points in **c**, whose abundance variation is attributed predominantly to temporal (red) or spatial sources (blue). Gray lines correspond to abundances of technical replicates (X,Y) obtained from the same spatial location, and colored lines correspond to abundances from the second spatial replicate (Z).

**Figure 3.**
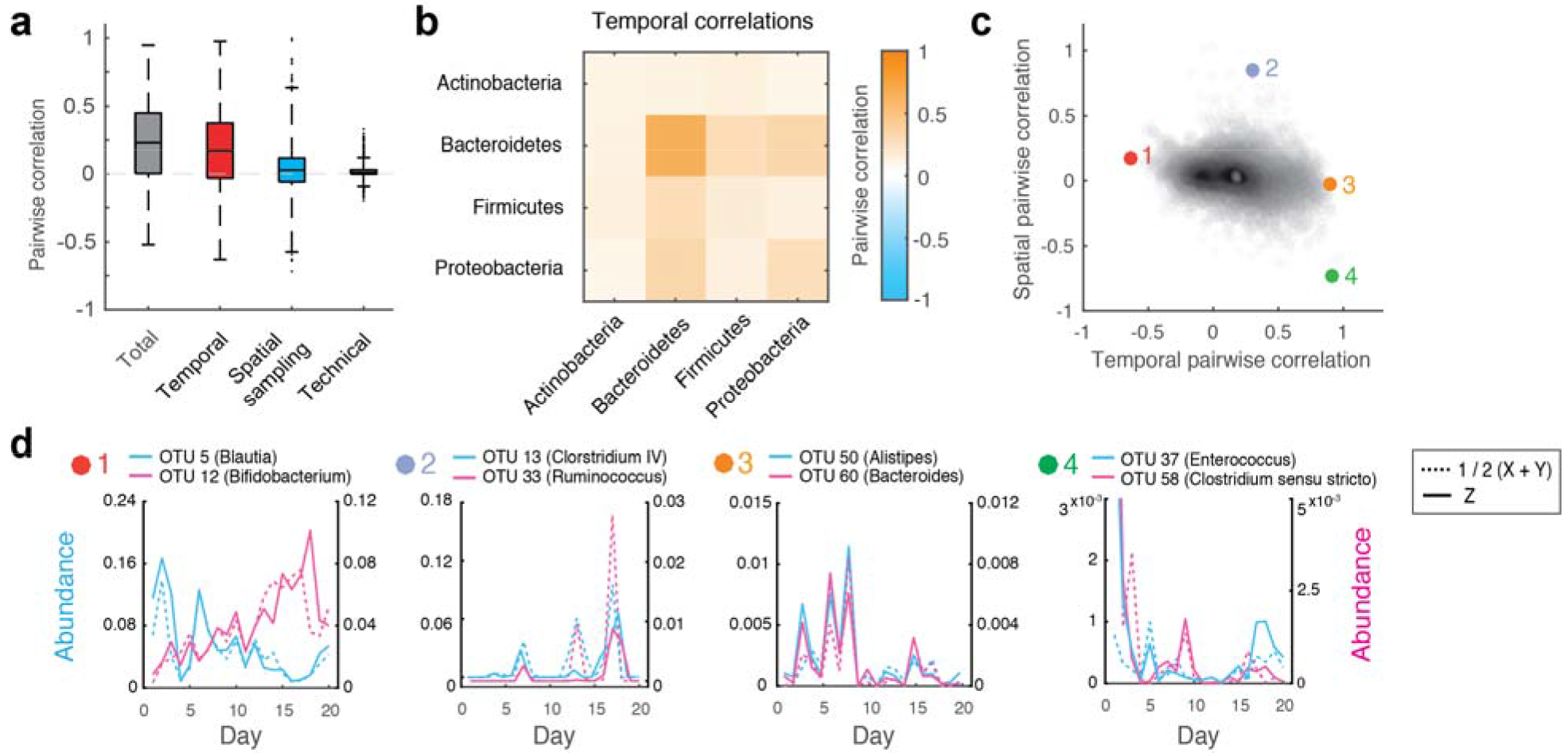
Decomposition of pairwise OTU abundance correlations in the human gut microbiome. (**a**) Boxplots of total, temporal, spatial and technical correlations for all pairs of abundant OTUs (average absolute abundance > 10^−4^). Boxes denote the median and interquartile ranges, with maximum whisker lengths three times the interquartile range. (**b**) Temporal correlations of OTU abundances within and between different phyla. Colors reflect average temporal correlations between pairs of OTUs belonging to the indicated phyla. Data are shown for all highly abundant OTUs (mean absolute abundance > 10^−4^) from the *Actinobacteria* (n = 10), *Bacteroidetes* (n=15), *Firmicutes* (n=103), and *Proteobacteria* (n=5). (**c**) Temporal and spatial correlations for all pairs of abundant OTUs (average absolute abundance > 10^−4^). Colored points (1-4) indicate pairs of OTUs with temporal profiles shown in **d.** (**d**) Temporal abundance profiles for pairs of OTUs highlighted in **c**. Pairs (from left to right) exhibit: 1) Substantial negative temporal (*ρ*_*T*_ = −0.63), 2) substantial positive spatial (*ρ*_*S*_ = 0.85), 3) substantial positive temporal (*ρ*_*T*_ = 0.90), and 4) substantial negative spatial (*ρ*_*S*_ = −0.73) correlations. For every OTU pair, blue and pink solid lines show abundances of each OTU measured from the same spatial location (Z). Blue and pink dashed lines show the average between technical replicates (1/2(X+Y)) of each OTU measured from the second spatial location. Note that the large spatial correlation between OTUs 13 and 33 (panel 2) is reflected in similar profiles of the two dashed lines, as well as the two solid lines (measurements from the same spatial location); the lower temporal correlation between these OTUs is reflected in more dissimilar profiles of solid and dashed lines of the opposite color (measurements from different spatial locations). Also note that OTUs 37 and 58 exhibit anti-correlated spatial behavior despite abundances being correlated over time.

To validate the developed workflow and the variance decomposition model, we performed a set of controlled experiments that specifically eliminated either temporal or spatial variability from collected fecal samples (Online Methods). First, we obtained fecal samples from ten independent spatial locations of the same stool specimen. This effectively simulated five consecutive time points of the DIVERS protocol, but without any temporal contribution to microbiota variability. Second, to remove spatial variability, we carried out eight consecutive days of sampling with spatial replicates homogenized on each day before sequencing (Online Methods). Reassuringly, the model correctly predicted no temporal or spatial contributions to OTU abundance variability when the corresponding signals were absent from the data (**Fig. 1e**).

The ability of DIVERS to unmask temporal and spatial variances of individual OTUs makes it possible to use absolute bacterial abundances to investigate microbiota fluctuations in the human gut^20^. Interestingly, temporal variances calculated from DIVERS showed a robust power law dependence on average OTU abundance, following a relationship known as Taylor’s law in ecology^21^ (**Fig. 2a**, power law exponent *b* = 1.9, Pearson’s r = 0.97 on log-transformed means and variances). In addition, DIVERS allowed us to investigate the relationship between average absolute abundances and their spatial variances, which could be also well-described by a power law (**Fig. 2b**, *b* = 1.7, r = 0.96). These results suggest that, in contrast to the null model of randomly distributed OTU abundances across time and space (*b* = 1), bacterial species within the gut microbiome display significantly more complex spatiotemporal dynamics^9,22^.

Beyond general ecological relationships, we used our approach to identify specific taxa with particularly high spatial or temporal contributions to their total abundance variance (**Fig. 2c**, **Supplementary Table 3**, Online Methods). Interestingly, the time series of several high-abundance OTUs showed behavior primarily shaped by either spatial (OTU 13, Genus: Clostridium IV and OTU 122, Genus: Terrisporobacter) or temporal factors (OTU 12, Genus: Bifidobacterium and OTU 25, Genus: Lachnospiracea incertae sedis) (**Fig. 2d**). These examples demonstrate that DIVERS may be used to characterize the spatiotemporal dynamics of individual OTUs in the human gut.

Fluctuations in bacterial abundances often result from the collective behavior of multiple different taxa, whose interactions are reflected in correlated abundance changes^23^. DIVERS can also be used to quantify the factors contributing to abundance correlations between pairs of OTUs in a microbial community (Online Methods and **Supplementary Note**). Applying this analysis to the human gut microbiome, we found that the majority of pairwise abundance correlations were due to temporal sources, with relatively smaller contributions from spatial sampling location and technical noise (**Fig. 3a** and **Supplementary Fig. 4**). Consistent with previous results^13^, we also found that total correlations based on absolute abundance measurements were generally larger than correlations calculated using relative abundances, an effect primarily caused by the variance in total bacterial loads across samples (**Supplementary Fig. 5** and **Supplementary Note**).

Next, we examined factors contributing to the correlations of OTU abundances within and between the four most prevalent bacterial phyla in the human gut. Interestingly, the *Bacteroidetes* exhibited significantly larger intra-phyla temporal abundance correlations compared to the rest of the community (p < 1e−10, Wilcoxon rank sum test) (**Fig. 3b**, **Supplementary Fig. 6, Supplementary Fig. 7**). This result was also observed at the family level, and was not due to differences in 16S rRNA sequence similarity across taxa (**Supplementary Fig. 8** and **Supplementary Fig. 9**). The coordinated temporal changes of *Bacteroidetes* in the human gut may reflect fluctuations in the availability of dietary polysaccharides on each day that are specifically metabolized by these bacteria^24,25^, as well as previously observed cross-feeding interactions between these taxa^26,27^. In addition, our analysis revealed several interesting examples of OTU pairs with positive and negative correlation contributions from both temporal and spatial factors (**Fig. 3c,d**). These examples highlight the diversity of bacterial dynamics in the gut, and demonstrate the ability of DIVERS to disentangle the factors contributing to abundance correlations between different taxa.

While current sequencing technologies enable bacterial communities to be profiled at high temporal resolution, novel approaches are required to reveal key features of ecosystem dynamics. Our results demonstrate the ability of DIVERS to quantify both the spatiotemporal dynamics and noise profiles of microbial communities, while requiring only a small number of additional samples compared to current metagenomic sequencing protocols. Although we focus on human gut microbiome dynamics in this study, DIVERS can be readily applied to explore patterns of variation in any bacterial ecosystem across different hosts and environments (**Supplementary Note**). Moreover, given the flexibility of the developed quantitative framework, it can be easily extended to other sequencing-based applications, such as the characterization of human immune cell repertoires^28^ and gene expression changes in tumors^29^.

## Author contributions

B.W.J. and R.U.S. conceived the study, designed the data collection workflow and performed all data analysis. B.W.J. and P.D.D. developed the variance and covariance decomposition models. R.U.S. performed all experiments. H.W.W. and D.V. oversaw the project, and guided experiments and data analysis. All authors wrote the manuscript.

## Methods

### Ethical review

This study was approved and conducted under Columbia University Medical Center Institutional Review Board protocol AAAR0753. Written informed consent was obtained from the subject in the study, a healthy male adult.

### Sample collection and storage

Fecal samples were collected daily over the course of twenty days, with two additional samples taken on days 27 and 48 of the study. After defecation, inverted sterile 200 µL pipette tips (Rainin RT-L200F) were used to core out a small sample from the stool, and placed immediately in a sterile cryovial (Sarstedt 72.694.106). On each day of stool collection, two samples were obtained from independent spatial locations at least >5cm in distance using the same stool specimen. Samples were immediately placed in a −20 °C freezer and transferred to a −80 °C freezer for long term storage.

### Spike-in strain for calculation of bacterial absolute abundances

A spike-in approach was utilized during sample processing to allow for calculation of bacterial absolute abundance per mass of fecal matter. Sporocarcina pasteurii (ATCC 11859), an environmental bacterium not found in human feces, was grown to saturation in NH4-YE medium (ATCC medium 1376). It was then concentrated by centrifugation, resuspended in ~0.1X volume phosphate buffered saline with 20% glycerol, and stored in cryovials at −80 °C for subsequent use during genomic DNA extraction.

### Replicate fecal sampling experimental protocol

To enable decomposition of gut bacterial abundance variability into temporal, spatial and technical contributions, we utilized a replicate sampling approach. Specifically, on each day, two fecal samples were collected from random spatial locations on the same stool specimen. For one of these samples, two technical replicates were prepared in parallel by splitting the individual fecal core. Thus, a total of three samples were processed for each day of the time series: two technical replicates from a single spatial location (denoted samples X and Y) and a second spatial replicate (denoted sample Z). To further characterize technical noise, a single fecal sample was subjected to 12 independent rounds of sample processing and sequencing. Metadata associated with all samples are given in **Supplementary Table 1**. Theoretical details associated with the DIVERS approach are described in the **Supplementary Note**.

### Sample genomic DNA extraction

Genomic DNA (gDNA) extraction was performed using a custom liquid handling protocol based on the Qiagen MagAttract PowerMicrobiome DNA/RNA Kit (Qiagen 27500-4-EP) adapted for lower volumes. Briefly, a 96 well plate (Axygen P-DW-20-C) was loaded with 1 mL of 0.1 mm Zirconia Silica beads (Biospec 11079101Z) using a loading device (Biospec 702L). During sample processing, appropriate negative controls were run on each plate (i.e. water control). 10 uL of thawed and concentrated spike-in strain was added to each well. 10-100 mg of each sample (average 45.9 mg, standard deviation 14.7 mg) was added to the plate using a sterile plastic spatula, and the weight added for each sample was determined via an analytical balance. 750 µL of lysis solution was then added to each well (90 mL master mix, 9 mL 1M Tris HCl pH 7.5, 9 mL 0.5M EDTA pH 8.0, 11.25 mL 10% SDS, 22.5 mL Qiagen lysis reagent, 38.25 mL nuclease free water). The plate was centrifuged down for 1 min at 4500xg and a bead sealing mat was affixed to the plate (Axygen AM-2ML-RD). The plate was then placed on a bead beater (Biospec 1001) and subjected to bead beating for 5 min followed by 10 min for cooling. This bead beating cycle was repeated, for a total of 10 min of bead beating. The plate was centrifuged down for 5 min at 4500xg and 200uL of supernatant was transferred to a V-bottom microplate. 35 µL of Qiagen inhibitor removal solution was added to each well and mixed by vortexing, incubated 4 °C for 5 min, and the plate was again centrifuged down for 5 min at 4500xg. 100 µL of supernatant was removed from the plate and placed in a round-bottom plate (Corning 3795). The plate was then placed on a robotic liquid handler (Biomek 4000) for magnetic bead purification of the supernatant per the manufacturers recommendations but at a scaled volume; magnetic beads in binding solution were mixed in each well, and subjected to 3 washes with wash solution and elution in 100 uL of nuclease free water into a new plate.

### 16S rRNA amplicon sequencing

16S sequencing of the V4 region was performed utilizing a custom protocol and a dual indexing scheme adapted from Kozich et al^1^. Briefly, dual indexing sequencing primers were adapted from the previous study, but we utilized Illumina Nextera barcode sequences and altered 16S primers to match updated 505f and 806rB primers (see Table S2 for sequences). A 20 µL PCR amplification was set up in a 96 well skirted PCR microplate: 1 µM forward 5XX barcoded primer, 1 µM reverse 7XX barcoded primer, 1 µL prepared gDNA, 10 uL NEBNext Q5 Hot Start HiFi Master Mix (NEB M0543L), 0.2X final concentration SYBR Green I. A quantiative PCR amplification (98°C 30s; cycle: 98°C 20s, 55°C 20s, 65°C 60s, 65°C 5m) was performed and cycling was stopped during exponential amplification (typically 12-20 cycles) and the reaction was advanced to the final extension step.

The resulting PCRs were quantified utilizing a SYBR Green I dsDNA assay; 2 µL of PCR product was added to 198 µL of TE with 1X final concentration SYBR Green I and fluorescence was quantified on a microplate reader. Samples were pooled based on this quantification on a robotic liquid handler (Biomek 4000). The resulting ~390 bp amplicon from the pool was then gel-purified utilizing a 2% E-gel (Invitrogen) and Wizard SV gel extraction kit.

Final libraries were then quantified by Qubit dsDNA HS assay and sequenced on the Illumina MiSeq platform (V2 500 cycle kit) according to the manufacturers instructions with modifications. Specifically, the library was loaded at 10 pM with 20% PhiX spike-in, and custom sequencing primers were spiked into the MiSeq reagent cartridge (6 uL of 100 µM stock; well 12: read1, well 13: index1, well 14: read2).

### Sequence analysis and OTU clustering

Resulting sequence data was analyzed with the USEARCH^30^ pipeline. Specifically, raw reads were merged using the –fastq_mergepairs command with options –fastq_maxdiffs 10 –fastq_maxdiffpct 10. Merged sequences were filtered using the –fastq_filter command with options –fastq_maxee 1.0 and –fastq_minlen 240. Resulting sequences were dereplicated (–derep_fulllength), clustered into OTUs (–cluster_otus) and the merged reads were searched against OTUs sequences (–usearch_global) at 97% identity. Taxonomic assignments of OTUs were made using the RDP classifier^31^.

### Calculation of OTU absolute abundances

Total bacterial loads in each sample were calculated using the following formula:

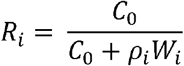

where, *R*_*i*_ is the sequenced relative abundance of the spike-in strain in sample *i*, *C*_*0*_ is the constant amount of spike-in strain (units of total DNA copies) added to each sample, *W_i_* is the weight of the fecal sample *i* (mg), and *ρ_i_* is the total bacterial density per fecal mass (DNA copies/mg). Solving for *ρ_i_*

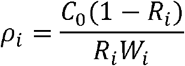

where we have measured *R*_*i*_ and *W*_*i*_ experimentally. Note that relative changes in *ρ*_*i*_ are independent of the constant *C*_0_. We therefore scaled total bacterial densities across samples to a mean of unity. Relative abundance profiles (with the spike-in strain excluded) were then multiplied by this scaled quantity to obtain absolute OTU abundances in arbitrary units that were used for all analyses.

### Variance decomposition of OTU abundances and total bacterial loads

DIVERS utilizes the replicate sampling and sequencing protocol described above to decompose measured bacterial abundance variances. Let *X* denote the total bacterial density in a collected sample or the abundance of an individual OTU. Using the law of total variance, the variance of *X* can be written as a sum of three components associated with temporal, spatial and technical factors contributing to changes in *X* across samples:

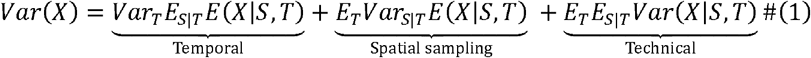

where, *S* and *T* are space and time-associated random variables capturing the spatial and temporal processes that influence the abundance of *X* across samples. Following the notation in **Fig. 1**, each of the terms in (1) is estimated as follows (see **Supplementary Note** for full derivations):

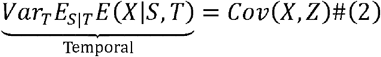

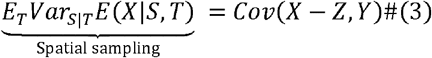

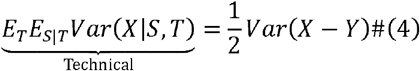

where *X, Z* and *Y, Z* denote pairs of spatial replicate measurements of either total bacterial density or individual OTU abundances. As described above, spatial replicates are obtained from two independent spatial locations in the environment at every time point. In contrast, *X* and *Y* denote technical replicates that are measured from the same spatial location.

### Covariance decomposition of OTU abundances

Using the law of total covariance, the covariance between the abundances of any two OTUs *i* and *j*, denoted *X*_*i*_ and *X*_*j*_, can also be written as a sum of temporal, spatial and technical contributions:

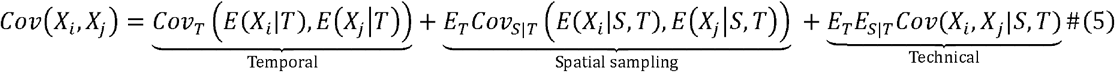

Each of the terms in (5) is estimated using the replicate sampling and sequencing protocol as follows (see **Supplementary Note** for full derivations):

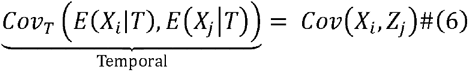

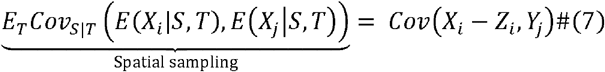

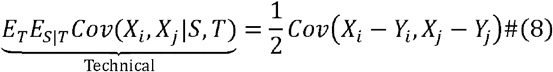

where, *X*_*i*_,*Z*_*i*_ and *Y*_*i*_,*Z*_*i*_ denote spatial replicate measurements of the abundance of OTU *i*, and *X*_*i*_,*Y*_*i*_ denote technical replicates. To obtain temporal, spatial and technical correlations shown in **Fig. 3**, we normalize each covariance contribution by the respective standard deviations of individual OTUs:

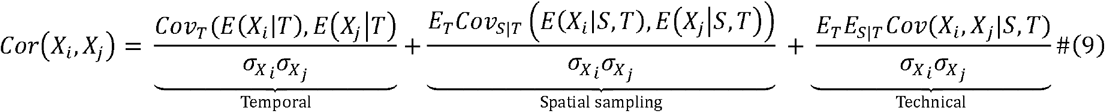

Variances and covariances of OTU abundances were calculated using data obtained across the twenty consecutive days of sampling. The variance decomposition of total bacterial densities also included samples taken from days 27 and 48 of the times series. To minimize artifacts due to technical noise, only OTUs with a mean absolute abundance >10^−4^ were included in the covariance decomposition analysis. This cutoff was chosen based on the observed variance profiles of individual OTUs (**Fig. 1d**). To compare contributions across phyla, 16S rRNA sequence-based phylogenetic distances were calculated using the pairwise2 module of Biopython.

### Identification of OTUs with high temporal or spatial variance contributions

To minimize effects of technical noise, OTUs were first filtered by abundance (mean absolute abundance >10^−4^). Of the remaining OTUs, those with temporal variance above 80% or spatial variance above 60% of total variability were identified and given in **Supplementary Table 3**.

### Removal of temporal or spatial variability from fecal samples

We conducted two sets of control experiments to remove either temporal or spatial variability of OTU abundances from fecal samples. Specifically, to eliminate temporal contributions, we re-sampled a single stool specimen ten times total to simulate five consecutive days of time series sampling. To eliminate spatial variability, replicate sampling was conducted for eight consecutive days; on each day, fecal samples obtained from random spatial locations were homogenized together by combining fecal samples, and then mechanically homogenizing in 1X phosphate buffered saline with a P200 pipette tip. The resulting homogenized sample was then split into technical triplicates and processed following the normal DIVERS protocol.

### Code availability

MATLAB scripts to perform all variance and covariance decomposition analyses from original OTU abundance tables will be available on GitHub at the time of publication.

### Data availability

All sequencing data will be made available through NCBI SRA at the time of publication.

