## Supplement Figures and Tables for "Quantifying spatiotemporal dynamics and noise in absolute microbiota abundances using replicate sampling"

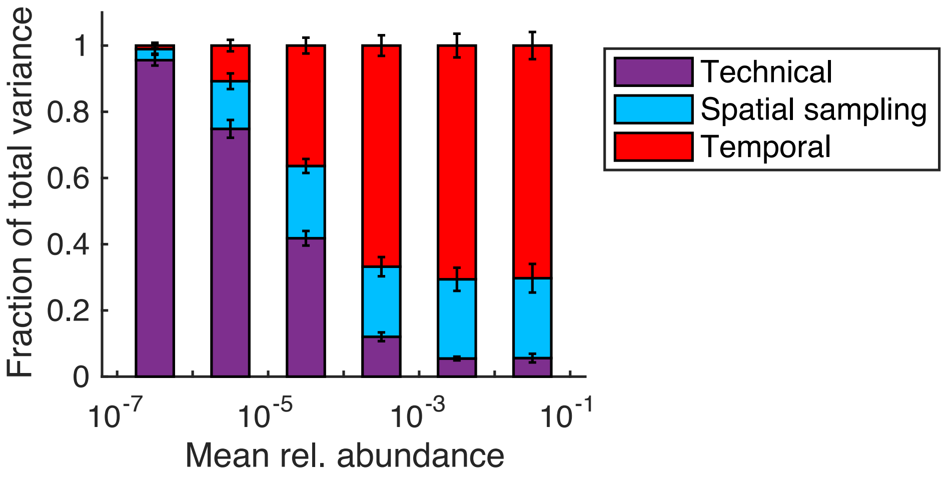


**Supplementary Figure 1**. Variance decomposition of OTU relative abundances. OTUs are binned by mean relative abundance across samples. Stacked bars indicate the average fraction of total variance attributed to temporal, spatial sampling and technical sources for OTUs within each bin. Error bars denote the SEM.

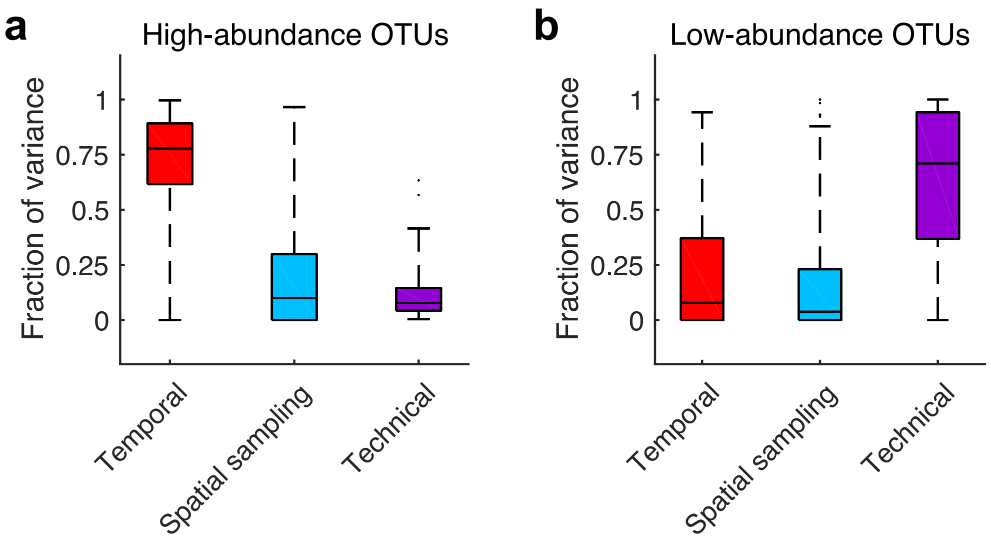


**Supplementary Figure 2**. Variance decomposition for high and low-abundance OTUs. Temporal, spatial sampling and technical contributions to total variance for all OTUs with (**a**) mean absolute abundance > $10^{-4}$ and (**b**) mean absolute abundance < $10^{-4}$. Boxes show the median and interquartile ranges, with maximum whisker lengths three times the interquartile range.

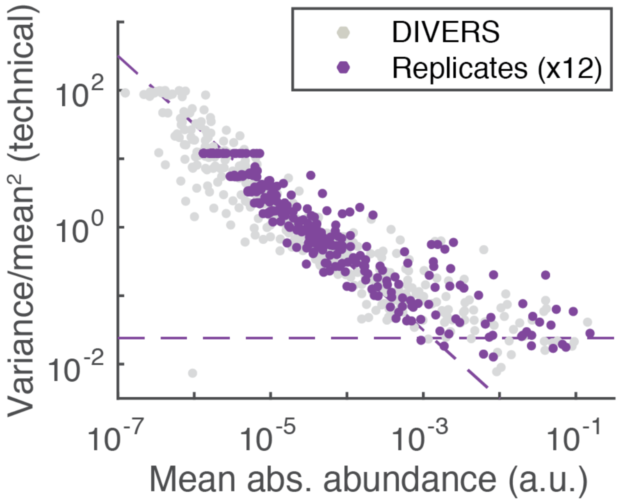


**Supplementary Figure 3.** OTU abundance variability due to technical noise. Multiple technical replicates (n=12) were processed from fecal samples obtained from a single spatial location of a stool specimen. Purple dots show the normalized technical variability (variance/mean$^2$) as a function of average abundance across twelve technical replicates. Technical noise profiles obtained from DIVERS are shown in gray dots. The inverse scaling expected from Poissonian sampling noise is indicated with the dashed line with slope = -1. A noise floor is observed at high OTU abundances (indicated by the horizontal dashed line) as a result of deviations in total bacterial load across samples.

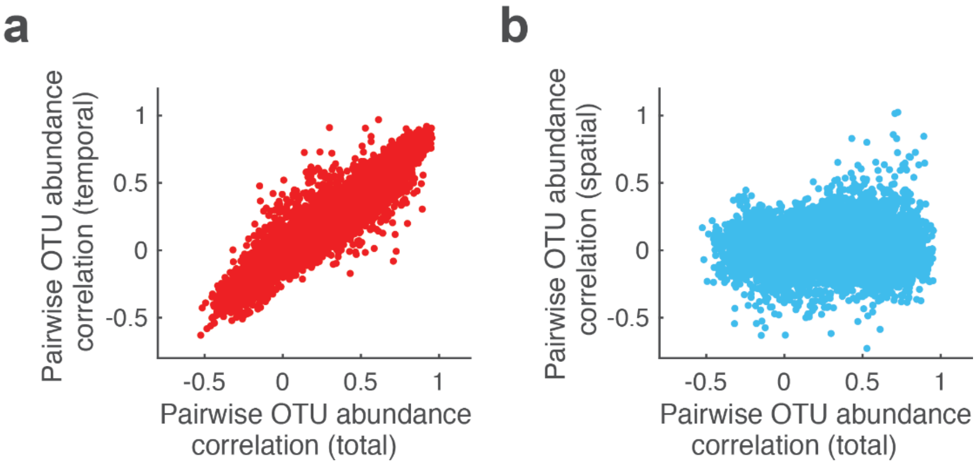


**Supplementary Figure 4**. Relationship between total pairwise OTU abundance correlations and (**a**) temporal and (**b**) spatial abundance correlations across all pairs of abundant OTUs (mean absolute abundance > $10^{-4}$). There is a significant correlation between temporal and total correlations (Pearson's r = 0.94, p < 1e-10).

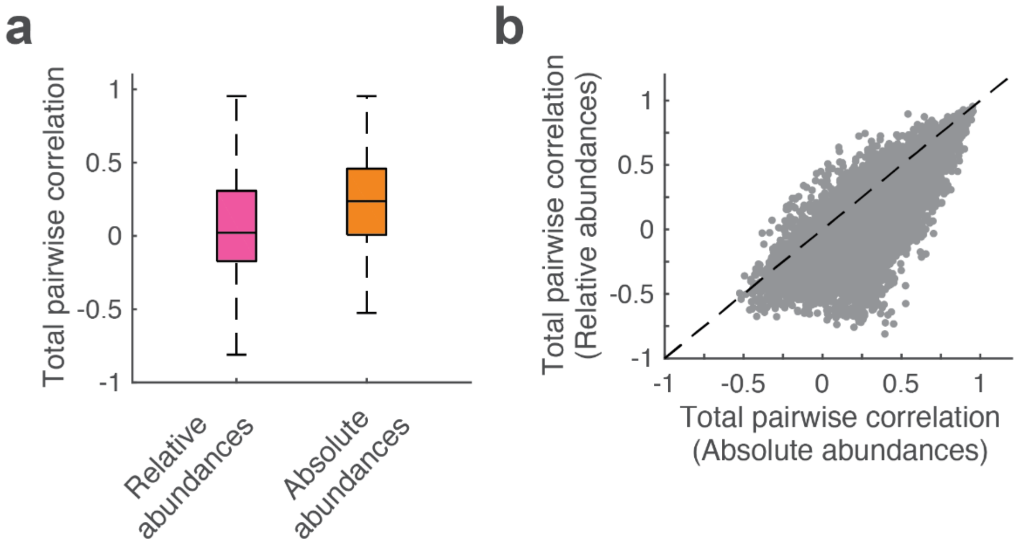


**Supplementary Figure 5**. Comparison between pairwise OTU abundance correlations calculated using relative or absolute abundances. (**a**) Total correlations calculated across all pairs of highly abundant OTUs (mean absolute abundance > $10^{-4}$) using relative (pink) or absolute (orange) abundances. Boxes show the median and interquartile ranges, with maximum whisker lengths three times the interquartile range. (**b**) Total pairwise OTU abundance correlations calculated using absolute abundances versus pairwise correlations using relative abundances; each point represents a pair of OTUs. Dashed line indicates the y = x line.

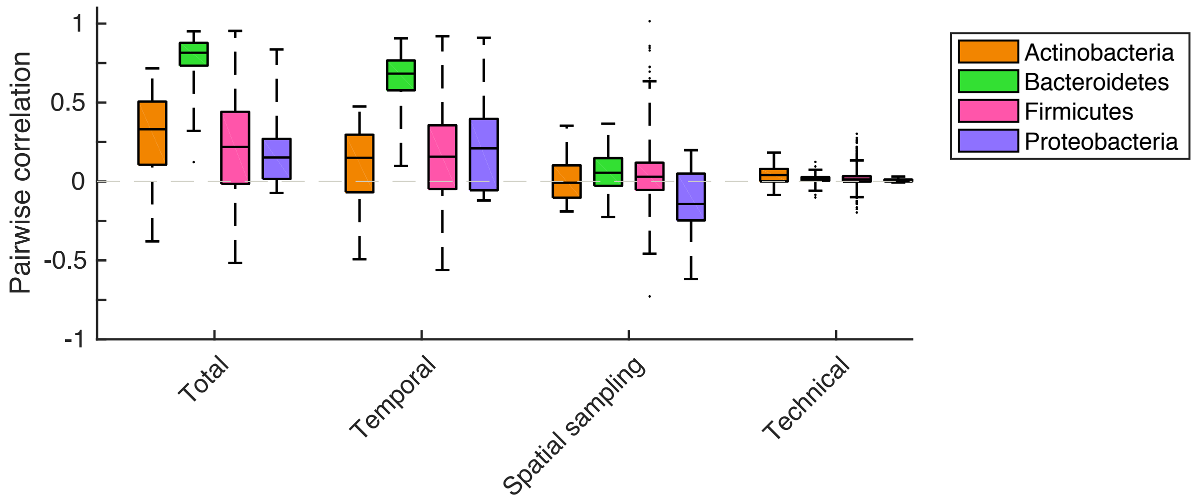


**Supplementary Figure 6.** Correlations of OTU abundances within different phyla. Boxplots indicate total, temporal, spatial and technical correlations of OTU abundances, where pairwise comparisons were made between OTUs within the indicated phyla. Boxes show median and interquartile ranges, with maximum whisker lengths three times the interquartile range.

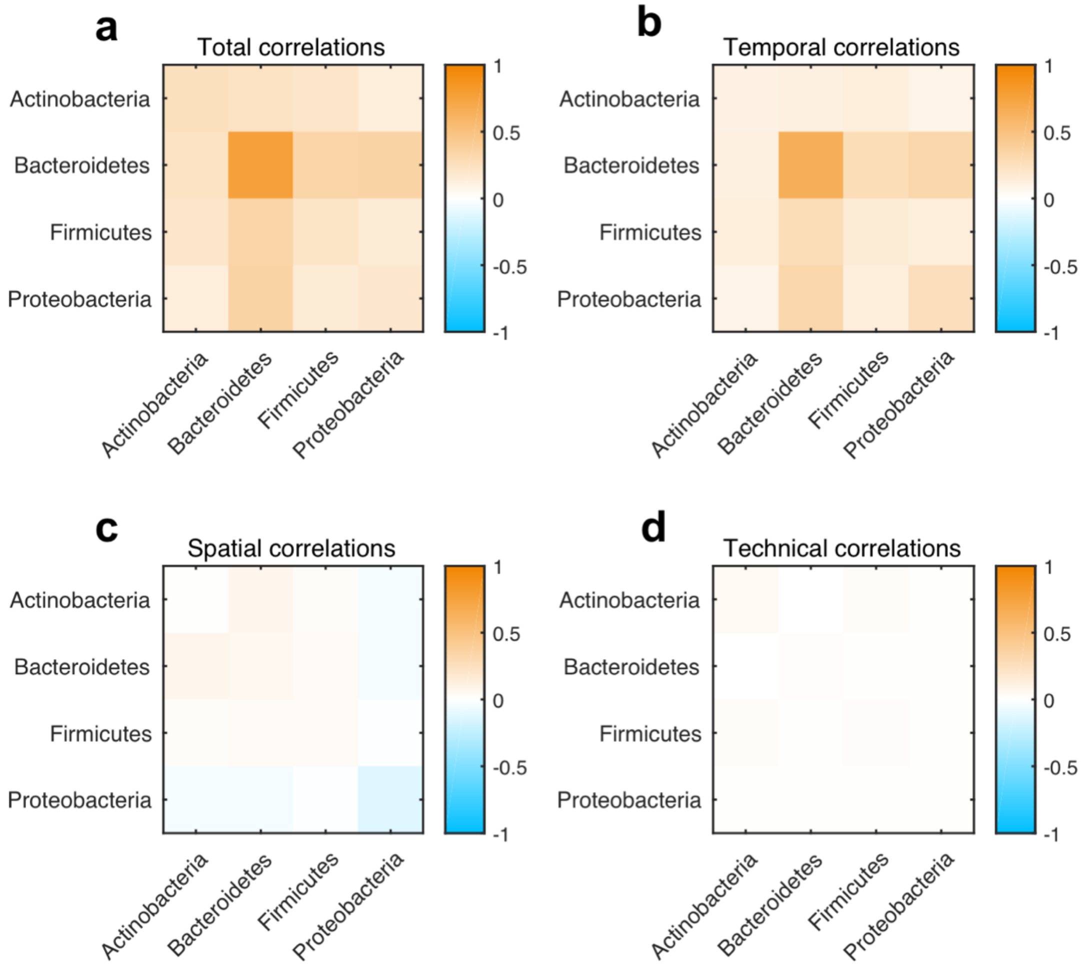


**Supplementary Figure 7.** Average correlations of OTU abundances within and between different phyla. Average (**a**) total, (**b**) temporal, (**c**) spatial and (**d**) technical correlations between pairs of OTUs belonging to the indicated phyla.

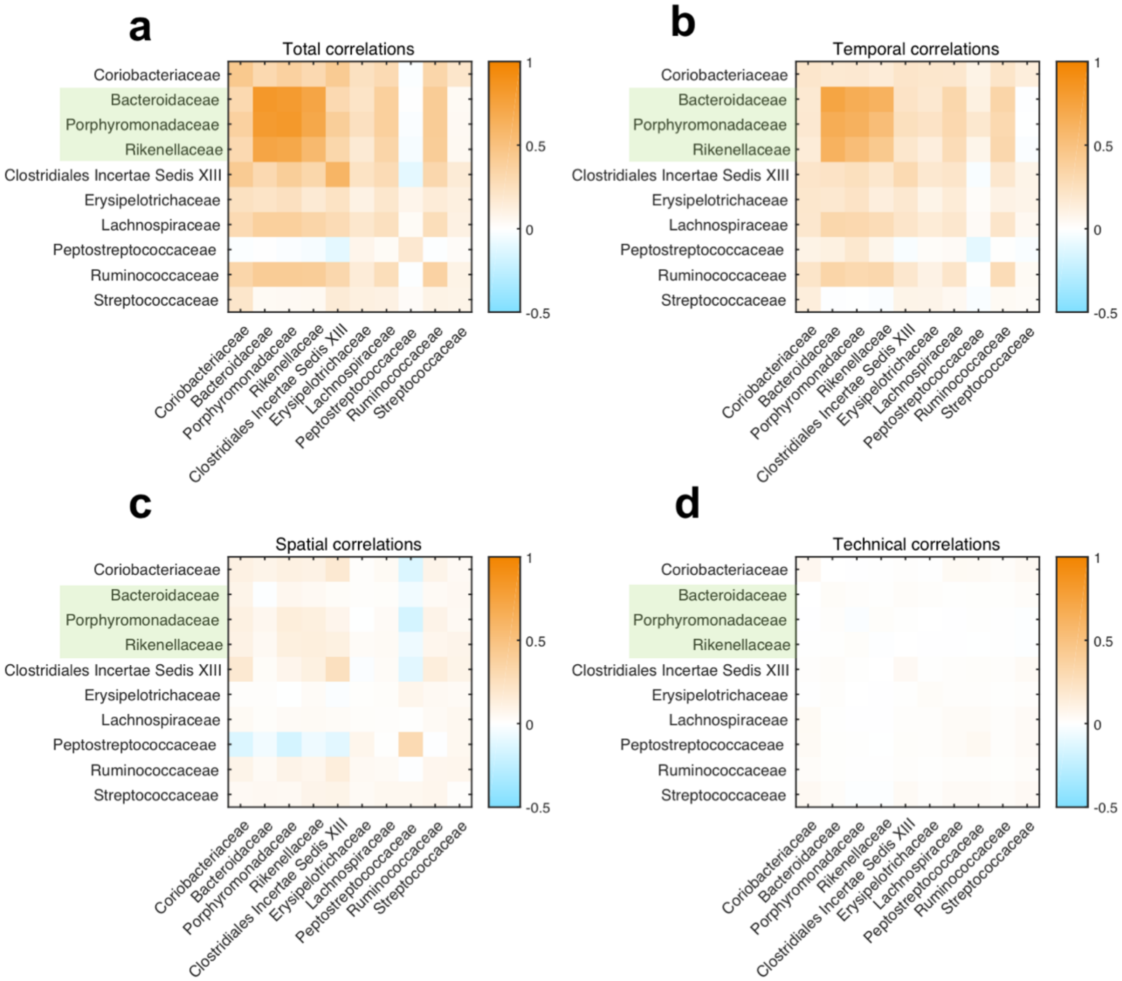


**Supplementary Figure 8.** Average correlations of OTU abundances within and between different microbial families. Average (**a**) total, (**b**) temporal, (**c**) spatial and (**d**) technical correlations between pairs of OTUs belonging to the indicated families. The three families belonging to the Bacteroidetes phylum are indicated with a green box.

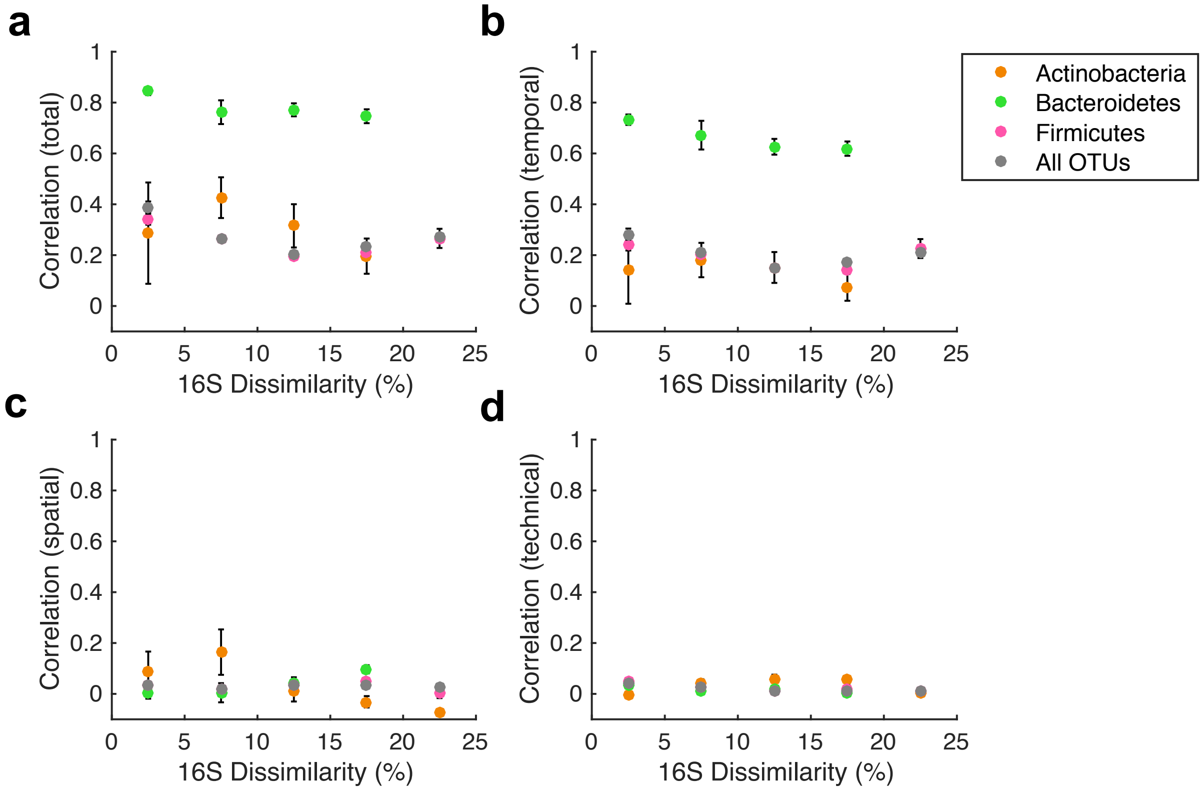


**Supplementary Figure 9.** Pairwise correlations of OTU abundances within phyla at different phylogenetic distances. (**a**) Total, (**b**) temporal, (**c**) spatial and (**d**) technical correlations between pairs of OTUs belonging to the same phyla were calculated for different bins representing the degree of 16S rRNA sequence dissimilarity. The mean correlations within each bin are shown for the indicated phyla with error bars denoting the SEM. Pairwise correlations for all abundant OTUs are shown in gray. Proteobacteria were excluded from the analysis due to insufficient sample size.

**Supplementary Table 1.** Fecal sample metadata for all samples analyzed in this study.

| Sample Name | Day | Time | Spatial_Rep. | Tech._Rep. | Weight (mg) |
|---|---|---|---|---|---|
| d16s1r1 | 1 | 9:00AM | 1 | 1 | 41.5 |
| d16s2r1 | 1 | 9:00AM | 2 | 1 | 85.5 |
| d16s2r2 | 1 | 9:00AM | 2 | 2 | 61.6 |
| d17s1r1 | 2 | 7:00PM | 1 | 1 | 49.5 |
| d17s2r1 | 2 | 7:00PM | 2 | 1 | 56.3 |
| d17s2r2 | 2 | 7:00PM | 2 | 2 | 52.2 |
| d18s1r1 | 3 | 3:00PM | 1 | 1 | 46.5 |
| d18s2r1 | 3 | 3:00PM | 2 | 1 | 63 |
| d18s2r2 | 3 | 3:00PM | 2 | 2 | 73.9 |
| d19s1r1 | 4 | 7:00PM | 1 | 1 | 57.4 |
| d19s2r1 | 4 | 7:00PM | 2 | 1 | 37.1 |
| d19s2r2 | 4 | 7:00PM | 2 | 2 | 37.5 |
| d20s1r1 | 5 | 12:00PM | 1 | 1 | 46.4 |
| d20s2r1 | 5 | 12:00PM | 2 | 1 | 39.6 |
| d20s2r2 | 5 | 12:00PM | 2 | 2 | 29.3 |
| d21s1r1 | 6 | 10:30PM | 1 | 1 | 49.9 |
| d21s2r1 | 6 | 10:30PM | 2 | 1 | 52.4 |
| d21s2r2 | 6 | 10:30PM | 2 | 2 | 41.5 |
| d22s1r1 | 7 | 11:56PM | 1 | 1 | 47.7 |
| d22s2r1 | 7 | 11:56PM | 2 | 1 | 36.7 |
| d22s2r2 | 7 | 11:56PM | 2 | 2 | 35.8 |
| d23s1r1 | 8 | 4:00PM | 1 | 1 | 43 |
| d23s2r1 | 8 | 4:00PM | 2 | 1 | 41 |
| d23s2r2 | 8 | 4:00PM | 2 | 2 | 44.4 |
| d24s1r1 | 9 | 4:30PM | 1 | 1 | 64.9 |
| d24s2r1 | 9 | 4:30PM | 2 | 1 | 38.8 |
| d24s2r2 | 9 | 4:30PM | 2 | 2 | 35.3 |
| d25s1r1 | 10 | 11:00PM | 1 | 1 | 59.2 |
| d25s2r1 | 10 | 11:00PM | 2 | 1 | 39.2 |
| d25s2r2 | 10 | 11:00PM | 2 | 2 | 56.4 |
| d26s1r1 | 11 | 10:00AM | 1 | 1 | 36.7 |
| d26s2r1 | 11 | 10:00AM | 2 | 1 | 58.8 |
| d26s2r2 | 11 | 10:00AM | 2 | 2 | 68.2 |
| d28s1r1 | 13 | 3:30PM | 1 | 1 | 57.4 |
| d28s2r1 | 13 | 3:30PM | 2 | 1 | 56.2 |
| d28s2r2 | 13 | 3:30PM | 2 | 2 | 45.5 |
| d29s1r1 | 14 | 3:00PM | 1 | 1 | 58 |
| d29s2r1 | 14 | 3:00PM | 2 | 1 | 53 |
| d29s2r2 | 14 | 3:00PM | 2 | 2 | 64.8 |
| d30s1r1 | 15 | 4:00PM | 1 | 1 | 71.4 |
| d30s2r1 | 15 | 4:00PM | 2 | 1 | 16.7 |
| d30s2r2 | 15 | 4:00PM | 2 | 2 | 23 |
| d31s1r1 | 16 | 8:30PM | 1 | 1 | 32.4 |
| d31s2r1 | 16 | 8:30PM | 2 | 1 | 63.5 |
| d31s2r2 | 16 | 8:30PM | 2 | 2 | 50.3 |
| d32s1r1 | 17 | 5:00PM | 1 | 1 | 62.3 |
| d32s2r1 | 17 | 5:00PM | 2 | 1 | 46.7 |
| d32s2r2 | 17 | 5:00PM | 2 | 2 | 68 |
| d33s1r1 | 18 | 12:45PM | 1 | 1 | 42.6 |
| d33s2r1 | 18 | 12:45PM | 2 | 1 | 38.1 |
| d33s2r2 | 18 | 12:45PM | 2 | 2 | 31.8 |
| d34as1r1 | 19a | 9:00AM | 1 | 1 | 41.7 |

| | | | | | |
|---|---|---|---|---|---|
| d34as2r1 | 19a | 9:00AM | 2 | 1 | 46.4 |
| d34as2r2 | 19a | 9:00AM | 2 | 2 | 35.2 |
| d34bs1r1 | 19b | 7:00PM | 1 | 1 | 57.1 |
| d34bs2r1 | 19b | 7:00PM | 2 | 1 | 46.3 |
| d34bs2r2 | 19b | 7:00PM | 2 | 2 | 46.2 |
| d35s1r1 | 20 | 4:00PM | 1 | 1 | 61.9 |
| d35s2r1 | 20 | 4:00PM | 2 | 1 | 44.6 |
| d35s2r2 | 20 | 4:00PM | 2 | 2 | 49 |
| d42s1r1 | 27 | 7:30PM | 1 | 1 | 58.3 |
| d42s2r1 | 27 | 7:30PM | 2 | 1 | 47.7 |
| d42s2r2 | 27 | 7:30PM | 2 | 2 | 54.9 |
| d63s1r1 | 48 | 10:00PM | 1 | 1 | 49.5 |
| d63s2r1 | 48 | 10:00PM | 2 | 1 | 38.2 |
| d63s2r2 | 48 | 10:00PM | 2 | 2 | 28.4 |
| d30s3r1 | 15 | 4:00PM | 3 | 1 | 60.8 |
| d30s4r1 | 15 | 4:00PM | 4 | 1 | 39.5 |
| d30s5r1 | 15 | 4:00PM | 5 | 1 | 44.7 |
| d30s6r1 | 15 | 4:00PM | 6 | 1 | 32.1 |
| d30s7r1 | 15 | 4:00PM | 7 | 1 | 54.1 |
| d30s8r1 | 15 | 4:00PM | 8 | 1 | 85.1 |
| d30s9r1 | 15 | 4:00PM | 9 | 1 | 39.7 |
| d30s10r1 | 15 | 4:00PM | 10 | 1 | 46.7 |
| d30s11r1 | 15 | 4:00PM | 11 | 1 | 65 |
| d30s12r1 | 15 | 4:00PM | 12 | 1 | 34.7 |
| d30s13r1 | 15 | 4:00PM | 13 | 1 | 49 |
| d30s14r1 | 15 | 4:00PM | 14 | 1 | 72.7 |
| d30s2r3 | 15 | 4:00PM | 2 | 3 | 26.4 |
| d30s2r4 | 15 | 4:00PM | 2 | 4 | 21.1 |
| d30s2r5 | 15 | 4:00PM | 2 | 5 | 29.1 |
| d30s2r6 | 15 | 4:00PM | 2 | 6 | 23.6 |
| d30s2r7 | 15 | 4:00PM | 2 | 7 | 15.6 |
| d30s2r8 | 15 | 4:00PM | 2 | 8 | 20.9 |
| d30s2r9 | 15 | 4:00PM | 2 | 9 | 11.8 |
| d30s2r10 | 15 | 4:00PM | 2 | 10 | 21 |
| d30s2r11 | 15 | 4:00PM | 2 | 11 | 16.4 |
| d30s2r12 | 15 | 4:00PM | 2 | 12 | 11.2 |

**Supplementary Table 2.** 16S sequencing primers utilized in the study.

| Primer Name | Sequence (5' to 3') |
|---|---|
| 16Sf_501 | AATGATACGGCGACCACCGAGATCTACAC TAGATCGC TATGGTAATT GT GTGYCAGCMGCCGCGGTAA |
| 16Sf_502 | AATGATACGGCGACCACCGAGATCTACAC CTCTCTAT TATGGTAATT GT GTGYCAGCMGCCGCGGTAA |
| 16Sf_503 | AATGATACGGCGACCACCGAGATCTACAC TATCCTCT TATGGTAATT GT GTGYCAGCMGCCGCGGTAA |
| 16Sf_504 | AATGATACGGCGACCACCGAGATCTACAC AGAGTAGA TATGGTAATT GT GTGYCAGCMGCCGCGGTAA |
| 16Sf_505 | AATGATACGGCGACCACCGAGATCTACAC GTAAGGAG TATGGTAATT GT GTGYCAGCMGCCGCGGTAA |
| 16Sf_506 | AATGATACGGCGACCACCGAGATCTACAC ACTGCATA TATGGTAATT GT GTGYCAGCMGCCGCGGTAA |
| 16Sf_507 | AATGATACGGCGACCACCGAGATCTACAC AAGGAGTA TATGGTAATT GT GTGYCAGCMGCCGCGGTAA |
| 16Sf_508 | AATGATACGGCGACCACCGAGATCTACAC CTAAGCCT TATGGTAATT GT GTGYCAGCMGCCGCGGTAA |
| 16Sr_701 | CAAGCAGAAGACGGCATACGAGAT TCGCCTTA AGTCAGTCAG CC GGACTACNVGGGTWTCTAAT |
| 16Sr_702 | CAAGCAGAAGACGGCATACGAGAT CTAGTACG AGTCAGTCAG CC GGACTACNVGGGTWTCTAAT |
| 16Sr_703 | CAAGCAGAAGACGGCATACGAGAT TTCTGCCT AGTCAGTCAG CC GGACTACNVGGGTWTCTAAT |
| 16Sr_704 | CAAGCAGAAGACGGCATACGAGAT GCTCAGGA AGTCAGTCAG CC GGACTACNVGGGTWTCTAAT |
| 16Sr_705 | CAAGCAGAAGACGGCATACGAGAT AGGAGTCC AGTCAGTCAG CC GGACTACNVGGGTWTCTAAT |
| 16Sr_706 | CAAGCAGAAGACGGCATACGAGAT CATGCCTA AGTCAGTCAG CC GGACTACNVGGGTWTCTAAT |
| 16Sr_707 | CAAGCAGAAGACGGCATACGAGAT GTAGAGAG AGTCAGTCAG CC GGACTACNVGGGTWTCTAAT |
| 16Sr_708 | CAAGCAGAAGACGGCATACGAGAT CCTCTCTG AGTCAGTCAG CC GGACTACNVGGGTWTCTAAT |
| 16Sr_709 | CAAGCAGAAGACGGCATACGAGAT AGCGTAGC AGTCAGTCAG CC GGACTACNVGGGTWTCTAAT |
| 16Sr_710 | CAAGCAGAAGACGGCATACGAGAT CAGCCTCG AGTCAGTCAG CC GGACTACNVGGGTWTCTAAT |
| 16Sr_711 | CAAGCAGAAGACGGCATACGAGAT TGCCTCTT AGTCAGTCAG CC GGACTACNVGGGTWTCTAAT |
| 16Sr_712 | CAAGCAGAAGACGGCATACGAGAT TCCTCTAC AGTCAGTCAG CC GGACTACNVGGGTWTCTAAT |
| 16S_read1 | TATGGTAATTGTGTGYCAGCMGCCGCGGTAA |
| 16S_read2 | AGTCAGTCAGCCGGACTACNVGGGTWTCTAAT |
| 16S_index1 | ATTAGAWACCCBNGTAGTCCGGCTGACTGACT |

**Supplementary Table S3.** OTUs with > 60% spatial or > 80% temporal abundance variance relative to the total variance. OTUs with high spatial variance are shaded in gray. Confidence of taxonomic assignments are indicated in parentheses.

| OTU | Phylum | Class | Order | Family | Genus |
|---|---|---|---|---|---|
| 13 | Firmicutes (98%) | Clostridia (98%) | Clostridiales (98%) | Ruminococcaceae (94%) | Clostridium IV (68%) |
| 33 | Firmicutes (100%) | Clostridia (100%) | Clostridiales (100%) | Ruminococcaceae (100%) | Ruminococcus (100%) |
| 48 | Firmicutes (100%) | Clostridia (100%) | Clostridiales (100%) | Peptostreptococcaceae (100%) | Intestinibacter (88%) |
| 71 | Verrucomicrobia (100%) | Verrucomicrobiae (100%) | Verrucomicrobiales (100%) | Verrucomicrobiaceae (100%) | Akkermansia (100%) |
| 75 | Firmicutes (100%) | Erysipelotrichia (100%) | Erysipelotrichales (100%) | Erysipelotrichaceae (100%) | Turicibacter (100%) |
| 101 | Proteobacteria (100%) | Gammaproteobacteria (100%) | Pasteurellales (100%) | Pasteurellaceae (100%) | Haemophilus (69%) |
| 122 | Firmicutes (100%) | Clostridia (100%) | Clostridiales (100%) | Peptostreptococcaceae (100%) | Terrisporobacter (100%) |
| 6 | Firmicutes (100%) | Clostridia (100%) | Clostridiales (100%) | Ruminococcaceae (100%) | Faecalibacterium (93%) |
| 11 | Firmicutes (100%) | Clostridia (100%) | Clostridiales (100%) | Lachnospiraceae (100%) | Fusicatenibacter (96%) |
| 12 | Actinobacteria (100%) | Actinobacteria (100%) | Bifidobacteriales (100%) | Bifidobacteriaceae (100%) | Bifidobacterium (100%) |
| 14 | Firmicutes (100%) | Erysipelotrichia (100%) | Erysipelotrichales (100%) | Erysipelotrichaceae (100%) | Clostridium XVIII (75%) |
| 22 | Firmicutes (100%) | Clostridia (100%) | Clostridiales (100%) | Ruminococcaceae (100%) | Ruminococcus (100%) |
| 24 | Bacteroidetes (100%) | Bacteroidia (100%) | Bacteroidales (100%) | Bacteroidaceae (100%) | Bacteroides (100%) |
| 25 | Firmicutes (100%) | Clostridia (100%) | Clostridiales (100%) | Lachnospiraceae (100%) | Lachnospiracea_incertae_sedis (75%) |
| 27 | Firmicutes (100%) | Clostridia (100%) | Clostridiales (100%) | Peptostreptococcaceae (100%) | Romboutsia (96%) |
| 28 | Firmicutes (100%) | Clostridia (100%) | Clostridiales (100%) | Lachnospiraceae (100%) | Lachnospiracea_incertae_sedis (92%) |
| 34 | Firmicutes (100%) | Clostridia (100%) | Clostridiales (100%) | Lachnospiraceae (99%) | Lachnospiracea_incertae_sedis (68%) |
| 35 | Firmicutes (100%) | Clostridia (100%) | Clostridiales (100%) | Ruminococcaceae (100%) | Faecalibacterium (77%) |
| 36 | Proteobacteria (100%) | Gammaproteobacteria (100%) | Enterobacteriales (100%) | Enterobacteriaceae (100%) | Escherichia/Shigella (79%) |
| 37 | Firmicutes (100%) | Bacilli (100%) | Lactobacillales (100%) | Enterococcaceae (100%) | Enterococcus (100%) |
| 39 | Proteobacteria (100%) | Gammaproteobacteria (100%) | Enterobacteriales (100%) | Enterobacteriaceae (100%) | Salmonella (98%) |
| 41 | Bacteroidetes (100%) | Bacteroidia (100%) | Bacteroidales (100%) | Bacteroidaceae (100%) | Bacteroides (100%) |
| 43 | Firmicutes (100%) | Negativicutes (100%) | Selenomonadales (100%) | Acidaminococcaceae (100%) | Phascolarctobacterium (100%) |
| 44 | Bacteroidetes (100%) | Bacteroidia (100%) | Bacteroidales (100%) | Bacteroidaceae (100%) | Bacteroides (100%) |
| 46 | Firmicutes (98%) | Clostridia (98%) | Clostridiales (98%) | Ruminococcaceae (74%) | Flavonifractor (19%) |
| 50 | Bacteroidetes (100%) | Bacteroidia (100%) | Bacteroidales (100%) | Rikenellaceae (100%) | Alistipes (100%) |
| 52 | Firmicutes (96%) | Clostridia (95%) | Clostridiales (95%) | Ruminococcaceae (78%) | Anaerotruncus (39%) |
| 54 | Firmicutes (100%) | Clostridia (100%) | Clostridiales (100%) | Lachnospiraceae (95%) | Clostridium XlVa (33%) |
| 55 | Firmicutes (100%) | Clostridia (100%) | Clostridiales (100%) | Lachnospiraceae (100%) | Roseburia (54%) |
| 56 | Firmicutes (100%) | Clostridia (100%) | Clostridiales (100%) | Ruminococcaceae (100%) | Oscillibacter (70%) |
| 57 | Firmicutes (100%) | Bacilli (100%) | Lactobacillales (100%) | Streptococcaceae (100%) | Lactococcus (100%) |
| 59 | Firmicutes (100%) | Negativicutes (100%) | Selenomonadales (100%) | Acidaminococcaceae (100%) | Acidaminococcus (100%) |
| 60 | Bacteroidetes (100%) | Bacteroidia (100%) | Bacteroidales (100%) | Bacteroidaceae (100%) | Bacteroides (100%) |
| 61 | Firmicutes (100%) | Clostridia (100%) | Clostridiales (100%) | Lachnospiraceae (98%) | Coprococcus (25%) |
| 62 | Bacteroidetes (100%) | Bacteroidia (100%) | Bacteroidales (100%) | Bacteroidaceae (100%) | Bacteroides (100%) |
| 64 | Firmicutes (100%) | Clostridia (100%) | Clostridiales (100%) | Lachnospiraceae (100%) | Clostridium XlVa (44%) |
| 67 | Firmicutes (100%) | Clostridia (100%) | Clostridiales (100%) | Lachnospiraceae (99%) | Anaerosporobacter (21%) |
| 68 | Firmicutes (100%) | Clostridia (100%) | Clostridiales (100%) | Ruminococcaceae (100%) | Clostridium IV (65%) |
| 69 | Firmicutes (100%) | Clostridia (100%) | Clostridiales (100%) | Lachnospiraceae (100%) | Lachnospiracea_incertae_sedis (47%) |
| 70 | Firmicutes (100%) | Clostridia (100%) | Clostridiales (100%) | Lachnospiraceae (100%) | Fusicatenibacter (58%) |
| 73 | Firmicutes (99%) | Clostridia (98%) | Clostridiales (98%) | Clostridiales_Incertae Sedis XIII (63%) | Mogibacterium (45%) |
| 76 | Firmicutes (100%) | Erysipelotrichia (100%) | Erysipelotrichales (100%) | Erysipelotrichaceae (100%) | Clostridium XVIII (100%) |
| 79 | Firmicutes (100%) | Clostridia (100%) | Clostridiales (100%) | Lachnospiraceae (99%) | Lachnobacterium (49%) |

| | | | | | |
|---|---|---|---|---|---|
| 81 | Firmicutes (100%) | Clostridia (100%) | Clostridiales (100%) | Ruminococcaceae (100%) | Clostridium IV (100%) |
| 85 | Firmicutes (100%) | Erysipelotrichia (100%) | Erysipelotrichales (100%) | Erysipelotrichaceae (100%) | Coprobacillus (84%) |
| 88 | Bacteroidetes (100%) | Bacteroidia (100%) | Bacteroidales (100%) | Porphyromonadaceae (100%) | Parabacteroides (100%) |
| 89 | Firmicutes (98%) | Clostridia (98%) | Clostridiales (98%) | Ruminococcaceae (93%) | Clostridium IV (81%) |
| 90 | Proteobacteria (100%) | Betaproteobacteria (100%) | Burkholderiales (100%) | Sutterellaceae (100%) | Parasutterella (100%) |
| 94 | Firmicutes (100%) | Clostridia (100%) | Clostridiales (100%) | Ruminococcaceae (99%) | Flavonifractor (77%) |
| 97 | Firmicutes (100%) | Clostridia (100%) | Clostridiales (100%) | Lachnospiraceae (98%) | Dorea (46%) |
| 99 | Firmicutes (100%) | Clostridia (100%) | Clostridiales (100%) | Ruminococcaceae (85%) | Sporobacter (41%) |
| 103 | Firmicutes (100%) | Clostridia (100%) | Clostridiales (100%) | Lachnospiraceae (100%) | Clostridium XlVa (57%) |
| 106 | Firmicutes (100%) | Clostridia (100%) | Clostridiales (100%) | Ruminococcaceae (99%) | Oscillibacter (98%) |
| 108 | Bacteroidetes (100%) | Bacteroidia (100%) | Bacteroidales (100%) | Rikenellaceae (100%) | Alistipes (100%) |
| 109 | Firmicutes (100%) | Clostridia (100%) | Clostridiales (100%) | Ruminococcaceae (100%) | Flavonifractor (100%) |
| 112 | Firmicutes (100%) | Clostridia (100%) | Clostridiales (100%) | Lachnospiraceae (93%) | Butyrivibrio (24%) |
| 114 | Firmicutes (100%) | Clostridia (100%) | Clostridiales (100%) | Ruminococcaceae (100%) | Flavonifractor (67%) |
| 116 | Firmicutes (100%) | Negativicutes (100%) | Selenomonadales (100%) | Veillonellaceae (100%) | Veillonella (100%) |
| 120 | Firmicutes (95%) | Clostridia (93%) | Clostridiales (93%) | Lachnospiraceae (50%) | Clostridium XlVb (50%) |
| 130 | Bacteroidetes (100%) | Bacteroidia (100%) | Bacteroidales (100%) | Bacteroidaceae (100%) | Bacteroides (100%) |
| 136 | Firmicutes (100%) | Clostridia (100%) | Clostridiales (100%) | Lachnospiraceae (99%) | Anaerosporobacter (25%) |
| 140 | Firmicutes (100%) | Clostridia (100%) | Clostridiales (100%) | Ruminococcaceae (100%) | Sporobacter (78%) |
| 305 | Firmicutes (100%) | Clostridia (100%) | Clostridiales (100%) | Lachnospiraceae (100%) | Clostridium XlVa (32%) |
| 354 | Firmicutes (100%) | Clostridia (100%) | Clostridiales (100%) | Ruminococcaceae (100%) | Faecalibacterium (100%) |
| 374 | Firmicutes (100%) | Clostridia (100%) | Clostridiales (100%) | Lachnospiraceae (100%) | Ruminococcus2 (100%) |
| 392 | Actinobacteria (100%) | Actinobacteria (100%) | Bifidobacteriales (100%) | Bifidobacteriaceae (100%) | Bifidobacterium (95%) |
