## Supplemental Note for "Quantifying spatiotemporal dynamics and noise in absolute microbiota abundances using replicate sampling"

### **Supplementary Note**: Full description of variance and covariance decomposition models

#### 1 Variance decomposition model

##### 1.1 Overview

Let $X_i$ be a random variable denoting the abundance of a given bacterial operational taxonomic unit (OTU) $i$ measured from 16S rRNA sequencing. Although we focus on single OTU abundances, the following also applies for total bacterial abundances in each sample. We let the measured abundances of OTU $i$ collected at different time points of a time series study reflect draws from an underlying distribution $p(X_i)$, which we refer to as the marginal distribution of $X_i$. We assume that there are three contributions to the total observed variability of $X_i$. The first corresponds to any set of temporal factors that systematically change from one day to another to drive changes in OTU abundances over time. This may encompass environmental factors, interspecies interactions and competition, and neutral drift in abundances. The second contribution reflects heterogeneity in the abundance of bacteria across different spatial locations in a given environment. Notably, this spatial sampling variability is inherent to some time series studies such as those of the gut microbiome, where measurements from fecal samples collected at different time points necessarily come from different spatial locations. We take the spatial sampling variability to reflect

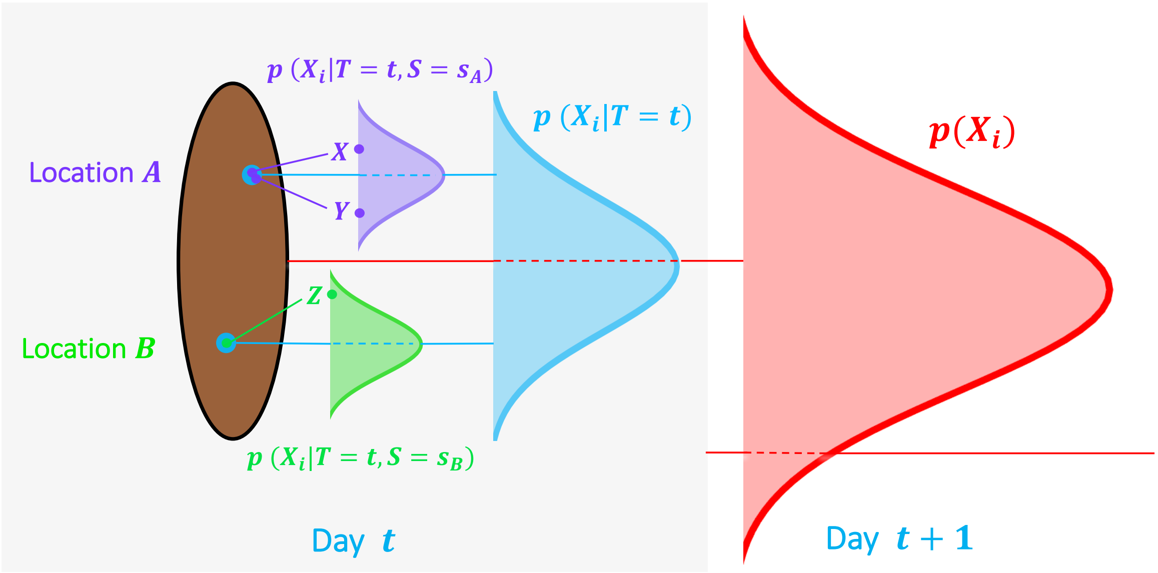

**Fig. N1**: Statistical model of the variance decomposition of bacterial abundances in the fecal microbiome.

variation arising from differences in niche size and availability or random dispersal of taxa abundances. The third corresponds to experimental noise associated with bacterial DNA extraction from samples, PCR amplification and sequencing itself. We collectively refer to these experimental sources of variability as technical noise. Our goal is to derive expressions for each of these sources of variability and demonstrate how one may estimate them from experiments.

#### 1.2 Decomposing the variance in microbiota abundances

We first demonstrate how we can use the law of total variance to decompose measured bacterial abundance variances into the three contributions described in the previous section.

##### *1.2.1 Defining the space and time variables*

As bacterial abundances change over time, we would expect that abundances of the same OTU $i$ measured at the same time point across different spatial locations in the environment would be more similar to each other than those collected from different time points. However, even at the same time point, measured abundances will not be identical due to both variation across different spatial locations and technical noise. Mathematically, the variability in abundances of OTU $i$ at a fixed time $T = t$ defines a conditional random variable $X_i^t$ with distribution $p(X_i|T = t)$, where $T$ is a time-associated random variable that captures the collective state of all time-associated factors and has underlying distribution $p(T)$. This conditional distribution itself may change when $T$ realizes different values at different time points. Importantly, however, when conditioned on a particular time point $T = t$, the variance of the conditional distribution $Var(X_i^t)$ reflects only spatial heterogeneity and technical noise.

At a given point in time, we may also choose a location from which to collect a sample to sequence. We can therefore define another random variable $X_i^{t,s}$ with probability distribution $p(X_i|T = t, S = s)$ representing the abundance of OTU $i$ measured from this fixed time point $T = t$ and spatial location $S = s$. Here, $S$ is a space-associated random variable with distribution $p(S|T = t)$ that at a given point in time, changes with the particular spatial location from which OTU $i$ is collected and measured. Conditioning on both space and time, we have eliminated any biological sources of variability and the variance of OTU $i$, $Var(X_i^{t,s})$, simply reflects technical noise. The distributions $p(X_i)$, $p(X_i|T = t)$, $p(X_i|T = t, S = s)$ and their hierarchical relationships are illustrated in Fig. N1, with the fecal microbiome shown as the model ecosystem.

##### 1.2.2 *Variance decomposition*

Using the law of total variance, we now decompose the total abundance variance of $X_i$ into components associated with time, spatial sampling location and technical noise. In the following, $E$ and $Var$ denote the expectation and variance of a random variable respectively, and subscripts denote the underlying distribution ($p(T)$ or $p(S|T)$) with respect to which the operation is performed. Beginning with the definition of the variance of $X_i$,

$$
\begin{aligned}
Var(X_i) &= E(X_i^2) - [E(X_i)]^2 \\
&= E_T E_{S|T} E(X_i^2|S,T) - [E_T E_{S|T} E(X_i|S,T)]^2 \\
&= E_T E_{S|T} Var(X_i|S,T) + E_T E_{S|T}[E(X_i|S,T)]^2 - [E_T E_{S|T} E(X_i|S,T)]^2 \\
&= E_T E_{S|T} Var(X_i|S,T) + E_T Var_{S|T} E(X_i|S,T) + E_T[E_{S|T} E(X_i|S,T|)]^2 - [E_T E_{S|T} E(X_i|S,T)]^2 \qquad (1)
\end{aligned}
$$

Thus,

$$
Var(X_i) = \underbrace{E_T E_{S|T} Var(X_i|S,T)}_{\text{Technical } (\langle\sigma_N^2\rangle_{S,T})} + \underbrace{E_T Var_{S|T} E(X_i|S,T)}_{\text{Spatial sampling } (\langle\sigma_S^2\rangle_T)} + \underbrace{Var_T E_{S|T} E(X_i|S,T)}_{\text{Temporal } (\sigma_T^2)} \qquad (2)
$$

The terms in the last line correspond to the contributions of technical, spatial sampling, and temporal factors to the total variance of $X_i$, which we denote with the symbols $\langle\sigma_N^2\rangle_{S,T}$, $\langle\sigma_S^2\rangle_T$ and $\sigma_T^2$ respectively. Equation (2) is simply the law of total variance generalized to multiple conditional random variables. Indeed, the right-most term can be recognized as the variance of $X_i$ explained by the time random variable $T$. The second term reflects the spatial sampling variance of OTU abundances conditioned on time, then averaged over time. The first term is simply the technical variability conditioned on a spatial location and time point, and then jointly averaged over space and time.

#### 1.3 Model-driven experimental approach

We now describe the approach we use to estimate the three contributions to total abundance variability derived in Section 1.2. We show mathematically how our model can be used to estimate each of these terms with only a minimal set of experiments. Notably, the approach we adopt is a generalization of the dual reporter method originally described by Elowtiz et. al. [1] used to separate intrinsic versus extrinsic sources of noise in the gene expression profiles of single cells [2, 3, 4].

##### 1.3.1 *Experimental setup*

Extending the notation described above, we denote $X_i$, $Y_i$ and $Z_i$ to be random variables representing abundances of an OTU $i$ made from three separate measurements at each point in time from the community. Let us assume that the abundances $X_i$ and $Y_i$ are made from the same exact location (by sequencing the sample twice), whereas $Z_i$ is measured from an independent location. As $X_i$, $Y_i$ and $Z_i$ correspond to the same bacterial OTU, their marginal distributions are equivalent. Importantly, however, because the abundances $X_i$,$Y_i$ and $Z_i$ are sampled together at the same time points, there exists a covariance structure driven by shared but unknown temporal factors tending to cause their abundances to collectively increase or decrease from one day to the next. Therefore, although $X_i$, $Y_i$ and $Z_i$ are identically distributed, they are not independent. In addition, by letting $X_i$ and $Y_i$ correspond to abundances measured not only at the same time point but also from the same spatial location, the covariance between $X_i$ and $Y_i$ is also driven by shared spatial factors that result in similar OTU abundances across different locations.

We can break the described covariance structures by conditioning abundances on time and spatial location. Specifically, when conditioning abundances on time $T = t$, the pairs $X_i^t, Z_i^t$ and $Y_i^t, Z_i^t$ have become sets of independent draws from the distribution $p(X_i|T = t)$. Experimentally, independence is achieved if $Z_i^t$ is sampled from a location in the community independent from $X_i^t$ and $Y_i^t$. Note that while $X_i^t$ and $Y_i^t$ have the same underlying distribution $p(X_i|T = t)$, they are not independent as their values covary across space. However, further conditioning of $X_i^t$ and $Y_i^t$ on spatial location results in the conditional random variables $X_i^{t,s}$ and $Y_i^{t,s}$ which are indeed independent draws from the distribution $p(X_i|T = t, S = s)$. The assumption of independence is reasonable, as $X_i^{t,s}$ and $Y_i^{t,s}$ are simply technical replicates.

Therefore, our replicate sampling protocol goes as follows: at each time point, we make three abundance measurements for all bacterial OTUs $i \in 1..N$. Two of these abundance measurements ($X_i$ and $Y_i$) are made from the same spatial location in a given environment. The third ($Z_i$) is measured from a separate, independent location. From an experimental standpoint, $X_i$ and $Y_i$ correspond to technical replicates while $X_i$,$Z_i$ and $Y_i$,$Z_i$ correspond to spatial replicates. Here, $t \in 1...n$ with $n$ being the total number of time points for which bacterial samples are collected.

#### 1.4 Derivation of statistical estimators for variance decomposition

We can now derive the statistical estimators for each of the terms in equation (2) using the complete hierarchical model described in Section 1.3. We begin by showing that under the specified model, the first two moments of the marginal distributions of $X_i$, $Y_i$, and $Z_i$ are indeed identical.

**Mean:** Using the law of total expectation,

$$E(X_i) = E_T E_{S|T} E(X_i|S, T) = E_T E_{S|T} E(Y_i|S, T) = E(Y_i) \tag{3}$$

$$E(X_i) = E_T E(X_i|T) = E_T E(Z_i|T) = E(Z_i) \tag{4}$$

**Variance:** By the law of total variance,

$$Var(X_i) = E_T E_{S|T} Var(X_i|S, T) + E_T Var_{S|T} E(X_i|S, T) + Var_T E_{S|T} E(X_i|S, T)$$

$$= E_T E_{S|T} Var(Y_i|S, T) + E_T Var_{S|T} E(Y_i|S, T) + Var_T E_{S|T} E(Y_i|S, T) = Var(Y_i) \tag{5}$$

$$Var(X_i) = E_T Var(X_i|T) + Var_T E(X_i|T) = E_T Var(Z_i|T) + Var_T E(Z_i|T) = Var(Z_i) \tag{6}$$

We have now laid the groundwork for the following derivations of statistical estimators for each of the terms in equation (2). This is the primary result of the variance decomposition model.

###### 1.4.1 Variance associated with time

$$\sigma_T^2 = Var_T E_{S|T} E(X_i|S,T) = Var_T E(X_i|T)$$

$$= E_T[E(X_i|T)]^2 - [E_T E(X_i|T)]^2$$

$$= E_T[E(X_i|T)E(Z_i|T)] - E_T E(X_i|T) E_T E(Z_i|T)$$

$$= E_T E(X_i Z_i|T) - E_T E(X_i|T) E_T E(Z_i|T)$$

$$= E(X_i Z_i) - E(X_i)E(Z_i) = Cov(X_i, Z_i) \tag{7}$$

Hence, the time-associated variability is simply the covariance between $X_i$ and $Z_i$, which can be easily estimated as:

$$\hat{\sigma_T^2} = \frac{1}{n-1}\sum_{t=1}^{n}(x_i^t - \bar{x_i})(z_i^t - \bar{z_i}) \tag{8}$$

###### 1.4.2 Variance associated with spatial sampling location

$$\langle\sigma_S^2\rangle_T = E_T Var_{S|T} E(X_i|S,T) = E_T E_{S|T}[E(X_i|S,T)]^2 - E_T[E_{S|T} E(X_i|S,T)]^2$$

$$= E_T E_{S|T}[E(X_i|S,T)]^2 - E_T[E(X_i|T)]^2$$

$$= E_T E_{S|T}[E(X_i|S,T)E(Y_i|S,T)] - E_T[E(Z_i|T)][E(Y_i|T)]$$

$$= E_T E_{S|T} E(X_i Y_i|S,T) - E_T E(Z_i Y_i|T)$$

$$= E(X_i Y_i) - E(Z_i Y_i)$$

$$= E(X_i Y_i - Z_i Y_i) - E(X_i)E(Y_i) + E(Z_i)E(Y_i)$$

$$= E((X_i - Z_i)Y_i) - E(X_i - Z_i)E(Y_i) = Cov(X_i - Z_i, Y_i) \tag{9}$$

Similar to time, the spatial sampling-associated variance reduces to the covariance between $X_i - Z_i$ and $Y_i$ and is estimated by:

$$\langle\hat{\sigma_S^2}\rangle_T = \frac{1}{n-1}\sum_{t=1}^{n}[(x_i^t - z_i^t) - (\bar{x_i} - \bar{z_i})](y_i^t - \bar{y_i}) \tag{10}$$

##### 1.4.3 Technical noise

$$\langle \sigma_N^2 \rangle_{S,T} = E_T E_{S|T} Var(X_i|S,T) = E_T E_{S|T} E(X_i^2|S,T) - E_T E_{S|T}[E(X_i|S,T)]^2$$

$$= \frac{1}{2}[E_T E_{S|T} E(X_i^2|S,T) - 2E_T E_{S|T}[E(X_i|S,T)E(Y_i|S,T)] + E_T E_{S|T} E(Y_i^2|S,T)]$$

$$= \frac{1}{2}[E(X_i^2) - 2E_T E_{S|T} E(X_i Y_i|S,T) + E(Y_i^2)]$$

$$= \frac{1}{2}[E(X_i^2) - 2E(X_i Y_i) + E(Y_i^2)]$$

$$= \frac{1}{2}[E(X_i^2) - 2E(X_i Y_i) + E(Y_i^2) - E(X_i)^2 + 2E(X_i)E(Y_i) - E(Y_i)^2]$$

$$= \frac{1}{2}[E([X_i - Y_i]^2) - [E(X_i - Y_i)]^2]$$

$$= \frac{1}{2} Var(X_i - Y_i) \tag{11}$$

Finally, the technical variability is simply the variance of the difference between $X_i$ and $Y_i$, which is estimated as:

$$\hat{\langle \sigma_N^2 \rangle}_{S,T} = \frac{1}{2(n-1)} \sum_{t=1}^{n} [(x_i^t - y_i^t) - (\bar{x}_i - \bar{y}_i)]^2 \tag{12}$$

#### 1.5 Generalizing the hierarchy

In the sections above, we have implicitly imposed a hierarchy in our model. Namely, quantities in each term of equation (2) are first averaged with respect to $p(X_i|S,T)$, followed by $p(S|T)$ and finally $p(T)$. Once can also imagine reversing this hierarchy; that is, averaging with respect to $p(T|S)$ second, followed by $p(S)$ last. Experimentally, this would correspond to drawing temporal replicates from a fixed spatial location, then averaging quantities over various locations. In ecosystems such as the human gut microbiome, the hierarchy described in earlier sections arises naturally, as $p(S|T)$ and $p(T)$ are experimentally accessible from fecal samples, while $p(T|S)$ and $p(S)$ are not. In other words, it is only possible to draw spatial replicates from a fixed time point, then average bacterial abundance measurements over multiple time points. It is not possible to measure abundances from the same spatial location on fecal samples obtained from two different days. However, in other microbial communities, one can imagine an alternative hierarchical sampling protocol in which different spatial locations in the ecosystem are each sampled on two separate days. Equation (2) may then be used to decompose bacterial abundance variances using this inverted hierarchy:

$$Var(X_i) = \underbrace{E_S E_{T|S} Var(X_i|S,T)}_{\text{Technical } (\langle \sigma_N^2 \rangle_{S,T})} + \underbrace{E_S Var_{T|S} E(X_i|S,T)}_{\text{Temporal } (\langle \sigma_T^2 \rangle_S)} + \underbrace{Var_S E_{T|S} E(X_i|S,T)}_{\text{Spatial sampling } (\sigma_S^2)} \tag{13}$$

The first term reflecting technical noise remains the same. The second term now reflects a temporal variance that is averaged over sampling locations. The third reflects the variance explained by spatial sampling location. We note that the spatial sampling variability here is defined rather loosely. For example, if one were to collect two

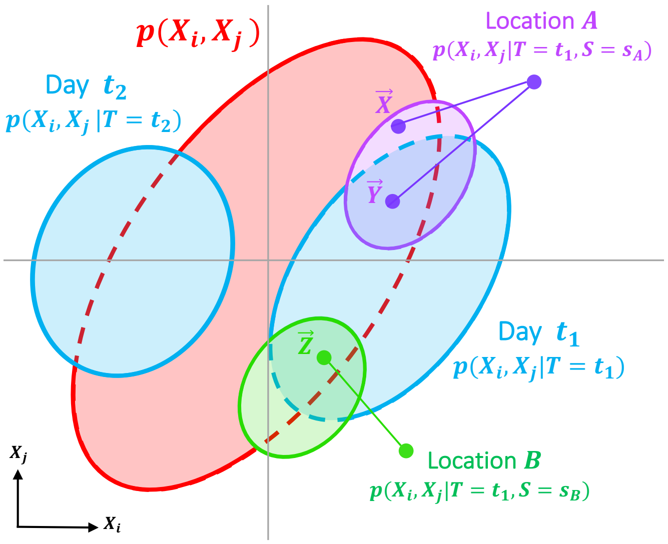

**Fig. N2**: Statistical model for bacterial abundance covariance decomposition.

human microbiome samples at different time points from a large cohort of individuals and sequence one of these two samples twice, the spatial sampling variability could be referred to as inter-individual variability while the temporal variability would correspond to intra-individual variability, with the noise term remaining the same. Finally, we remark that abundance measurements may be made from multiple different spatial locations at multiple different time points, with multiple sequencing replicates performed at each time and location. While experimentally much more demanding, one may arrive at the different variance contributions directly by following the prescription of equation (2) and using the typical estimators for mean and variance. However, our hierarchical replicate sampling protocol makes such data collection unnecessary.

### 2 Covariance decomposition model

#### 2.1 Overview

We now extend our variance decomposition model for single OTUs to a generalized covariance decomposition for all pairs of OTUs. We let $X_i$ and $X_j$ denote the abundances of OTUs $i$ and $j$ measured together from the same sample (i.e. sequenced from the same spatial location in a given environment). As with the case of single OTU variances, we assume that the total abundance covariance between OTUs $i$ and $j$ (across different samples collected over time) may be attributed to underlying temporal, spatial and technical sources. Intuitively, the temporal contribution results from the covariation in overall abundances (averaged over all spatial locations in the community) of OTUs $i$ and $j$ from one day to the next. In addition, however, species abundances may be correlated across different spatial locations at a given time point, which is captured by the spatial contribution to the total covariance. Finally, technical factors may result in correlated noise, potentially arising from sources such as similar DNA extraction efficiencies or primer and amplification biases. Again, our goal is to derive expressions for each of these covariance sources and demonstrate how one may estimate them experimentally using the same protocol described in Section 1.

#### 2.2 Decomposing the covariance in pairs of bacterial species abundances

##### *2.2.1 Total and conditional joint distributions*

The total abundance covariance of OTUs $i$ and $j$ may be calculated from the simple experiment: draw a single sample from a random spatial location in the environment at each time point and sequence the abundances of $i$ and $j$. Mathematically, we may consider the bivariate random variable $\vec{X}$ with components comprising the measured abundances $(X_i, X_j)$ and define a total joint distribution $p(X_i, X_j)$. We refer to this as the total joint distribution because in the data collection process, we have marginalized over time, space and technical noise. In contrast, one can imagine a second experiment where at a given time point, multiple samples are obtained across various spatial locations in the community. This defines another distribution, the conditional joint distribution $p(X_i, X_j|T = t)$ for some fixed time point $t$, where variances and covariances now reflect both underlying spatial factors as well as technical noise. Finally, by fixing both time and sampling location $S = s$ and re-sequencing the extracted DNA multiple times, a third distribution $p(X_i, X_j|T = t, S = s)$ can be defined. Here, variances and covariances reflect purely technical sources. The hierarchical relationships of these distributions are illustrated in Fig. N2. We show in the next section how the total covariance $Cov(X_i, X_j)$ between OTUs $i$ and $j$ may be decomposed by making use of the described conditional joint distributions.

###### 2.2.2 Covariance decomposition

The total covariance corresponding to the distribution $p(X_i, X_j)$ can be written as:

$$\begin{aligned}
Cov(X_i, X_j) &= E(X_iX_j) - E(X_i)E(X_j) \\
&= E_T E_{S|T} E(X_iX_j|S,T) - E(X_i)E(X_j) \\
&= E_T E_{S|T} Cov(X|S,T, Y|S,T) + E_T E_{S|T}[E(X_i|S,T)E(X_j|S,T)] - E(X_i)E(X_j) \\
&= E_T E_{S|T} Cov(X|S,T, Y|S,T) + E_T Cov_{S|T}(E(X_i|S,T), E(X_j|S,T)) \\
&\quad + E_T[E_{S|T}E(X_i|S,T)E_{S|T}E(X_j|S,T)] - E(X_i)E(X_j)
\end{aligned} \tag{14}$$

Mirroring the variance decomposition, we arrive at:

$$Cov(X_i, X_j) = \underbrace{E_T E_{S|T} Cov(X_i, X_j|S,T)}_{\text{Technical } (\langle \sigma^i \sigma^j{}_N \rangle_{S,T})} + \underbrace{E_T Cov_{S|T}(E(X_i|S,T), E(X_j|S,T))}_{\text{Spatial sampling } (\langle \sigma^i \sigma^j{}_S \rangle_T)} + \underbrace{Cov_T(E(X_i|T), E(X_j|T))}_{\text{Temporal } (\sigma^i \sigma^j{}_T)} \tag{15}$$

Note that we can obtain correlations by simply dividing each term in equation (15) by marginal standard deviations.

##### 2.3 Derivation of statistical estimators for covariance decomposition

Let $X_i, Y_i$ and $Z_i$ and $X_j, Y_j$ and $Z_j$ denote abundances of OTUs $i$ and $j$ respectively as described in Section 1.3.1. That is, the pairs $(X_i, Z_i)$ and $(Y_i, Z_i)$ correspond to spatial replicates while $(X_i, Y_i)$ denote technical replicates. As before, when conditioning on time $T = t$, the bivariate random variable pairs $\vec{X}^t = (X_i^t, X_j^t)$ and $\vec{Z}^t = (Z_i^t, Z_j^t)$, and $\vec{Y}^t = (Y_i^t, Y_j^t)$ and $\vec{Z}^t = (Z_i^t, Z_j^t)$ correspond to two i.i.d. draws from the same underlying distribution $p(X_i, X_j|T = t)$, with a covariance matrix structured by both spatial and technical sources. We will now define two additional and equivalent conditional joint distributions $p(X_i, Z_j|T = t)$, and $p(Y_i, Z_j|T = t)$ where the abundance of OTU $i$ from one spatial replicate $X_i^t$ or $Y_i^t$ is paired with the abundance of OTU $j$ from the second spatial replicate $Z_j^t$. Note here that while the conditional marginal distributions of $p(X_i, Z_j|T = t)$ and $p(Y_i, Z_j|T = t)$ are identical to those of the distribution $p(X_i, X_j|T = t)$ as demonstrated in Section 1.3, the random variables $X_i^t$ and $Z_j^t$, and $Y_i^t$ and $Z_j^t$ are independent of one another when conditioned on time, as abundances from each pair come from different spatial locations and sequencing realizations. Along similar lines, the bivariate random variables $\vec{X}^{t,s} = (X_i^{t,s}, X_j^{t,s})$ and $\vec{Y}^{t,s} = (Y_i^{t,s}, Y_j^{t,s})$ correspond to random variables drawn from the distribution $p(X_i, X_j|T = t, S = s)$, where correlations between $i$ and $j$ are driven purely by technical sources. Again, we may preserve conditional marginal distributions while eliminating covariances by defining the distribution $p(X_i, Y_j|T = t, S = s)$. With this in mind, we will now derive statistical estimators for each of the terms in equation (15).

###### 2.3.1 Covariance associated with time

$$
\begin{aligned}
\sigma^i\sigma^j{}_T &= Cov_T(E(X_i|T), E(X_j|T)) \\
&= E_T[E(X_i|T)E(X_j|T)] - E_TE(X_i|T)E_TE(X_j|T) \\
&= E_T[E(X_i|T)E(Z_j|T)] - E_TE(X_i|T)E_TE(Z_j|T) \\
&= E_TE(X_iZ_j|T) - E(X_i)E(Z_j) \\
&= Cov(X_i, Z_j)
\end{aligned} \tag{16}
$$

Analogous to the variance decomposition, the time-associated covariance can be easily estimated as:

$$
\hat{\sigma^i}\hat{\sigma^j}_T = \frac{1}{n-1}\sum_{t=1}^{n}(x_i^t - \bar{x_i})(z_j^t - \bar{z_j}) \tag{17}
$$

###### 2.3.2 Covariance associated with spatial sampling location

$$
\begin{aligned}
\langle\sigma^i\sigma^j{}_S\rangle_T &= E_TCov_{S|T}(E(X_i|S,T), E(X_j|S,T)) \\
&= E_TE_{S|T}[E(X_i|S,T)E(X_j|S,T)] - E_T[E_{S|T}E(X_i|S,T)E_{S|T}E(X_j|S,T)] \\
&= E_TE_{S|T}[E(X_i|S,T)E(X_j|S,T)] - E_T[E(X_i|T)E(X_j|T)] \\
&= E_TE_{S|T}[E(X_i|S,T)E(Y_j|S,T)] - E_T[E(Z_i|T)E(Y_j|T)] \\
&= E_TE_{S|T}E(X_iY_j|S,T) - E_TE(Z_iY_j|T) \\
&= E(X_iY_j) - E(Z_iY_j) \\
&= E(X_iY_j) - E(Z_iY_j) - E(X_i)E(Y_j) + E(Z_i)E(Y_j) \\
&= E((X_i - Z_j)Y_j) + E(X_i - Z_i)E(Y_j) \\
&= Cov(X_i - Z_i, Y_j)
\end{aligned} \tag{18}
$$

The space-associated covariance is thus given by:

$$
\langle\sigma^i\sigma^j{}_S\rangle_T = \frac{1}{n-1}\sum_{t=1}^{n}[(x_i^t - z_i^t) - (\bar{x_i} - \bar{z_i})](y_j^t - \bar{y_j}) \tag{19}
$$

###### 2.3.3 Covariance associated with technical noise

$$\langle \sigma^i \sigma^j{}_N \rangle_{S,T} = E_T E_{S|T} Cov(X_i, X_j|S,T)$$

$$= E_T E_{S|T} E(X_i X_j|S,T) - E_T E_{S|T}[E(X_i|S,T)E(X_j|S,T)]$$

$$= \frac{1}{2}[E_T E_{S|T} E(X_i X_j|S,T) - E_T E_{S|T}[E(X_i|S,T)E(Y_j|S,T)]$$
$$- E_T E_{S|T}[E(Y_i|S,T)E(X_j|S,T)] + E_T E_{S|T} E(Y_i Y_j|S,T)]$$

$$= \frac{1}{2}[E_T E_{S|T} E(X_i X_j|S,T) - E_T E_{S|T} E(X_i Y_j|S,T)$$
$$- E_T E_{S|T} E(Y_i X_j|S,T) + E_T E_{S|T} E(Y_i Y_j|S,T)]$$

$$= \frac{1}{2}[E(X_i X_j) - E(X_i Y_j) - E(Y_i X_j) + E(Y_i Y_j)]$$

$$= \frac{1}{2}[E(X_i X_j - X_i Y_j - Y_i X_j + Y_i Y_j) - E(X_i)E(X_j) + E(X_i)E(Y_j) - E(Y_i)E(X_j) + E(Y_i)E(Y_j)]$$

$$= \frac{1}{2}[E(X_i - Y_i)(X_j - Y_j) - E(X_i - Y_i)E(X_j - Y_j)]$$

$$= \frac{1}{2} Cov(X_i - Y_i, X_j - Y_j) \tag{20}$$

Finally, the covariance associated with technical sources is estimated as:

$$\langle \sigma^i \sigma^j{}_N \rangle_{S,T} = \frac{1}{2(n-1)} \sum_{t=1}^{n} [(x_i^t - y_i^t) - (\bar{x_i} - \bar{y_i})][(x_j^t - y_j^t) - (\bar{x_j} - \bar{y_j})] \tag{21}$$

###### 2.3.4 Covariances induced by conversion to absolute abundances

Finally, we demonstrate that conversion from relative to absolute abundances typically results in higher covariances for OTU pairs measured in absolute abundances. These higher covariances stem from the variance of total bacterial load measured from sample to sample that may induce correlations in OTU pairs whose relative abundances are otherwise uncorrelated or even negatively correlated. Let $R_i$ and $R_j$ denote the relative abundances of OTUs $i$ and $j$ measured from a single sample. Let us denote $A$ to be the total bacterial abundance density (in units of DNA copies per mg of environmental sample matter). Note that the absolute abundances $X_i$ and $X_j$ are simply calculated as $AR_i$ and $AR_j$ respectively. We will assume that the relative abundances $R_i$ and $R_j$ are independent of $A$. Then, we may write:

$$Cov(AR_i, AR_j) = E(A^2 R_i R_j) - E(A)^2 E(R_i)E(R_j)$$

$$= E(A^2)E(R_i R_j) - E(A)^2 E(R_i)E(R_j)$$

$$= E(A^2)[Cov(R_i, R_j) + E(R_i)E(R_j)] - E(A)^2 E(R_i)E(R_j)$$

$$= E(A^2)Cov(R_i, R_j) + E(R_i)E(R_j)[E(A^2) - E(A)^2)]$$

$$= E(A^2)Cov(R_i, R_j) + E(R_i)E(R_j)Var(A) \tag{22}$$
